# Spinal Microglia Contribute to Sustained Inflammatory Pain via Amplifying Neuronal Activity

**DOI:** 10.1101/2019.12.16.878728

**Authors:** Nan Gu, Min-Hee Yi, Madhuvika Murugan, Manling Xie, Jiyun Peng, Ukpong B Eyo, Hailong Dong, Long-Jun Wu

## Abstract

Microglia are highly dynamic immune cells of the central nervous system (CNS). Microglial processes interact with neuronal elements constantly on the order of minutes. The functional significance of this acute microglia-neuron interaction and its potential role in the context of pain is still largely unknown. Here, we found that spinal microglia increased their process motility and electrophysiological reactivity within an hour after the insult in a mouse model of formalin-induced acute, sustained, inflammatory pain. Using an ablation strategy to specifically deplete resident microglia in the CNS, we demonstrate that microglia participate in formalin-induced acute sustained pain behaviors by amplifying neuronal activity in the spinal dorsal horn. Moreover, we identified that the P2Y12 receptor, which is specifically expressed in microglia in the CNS, was required for microglial function in formalin-induced pain. Taken together, our study provides a novel insight into the contribution of microglia and the P2Y12 receptor in acute, sustained, inflammatory pain that could be used for potential therapeutic strategies.

## INTRODUCTION

Microglia are the resident immune cells of the central nervous system (CNS), which actively survey the microenvironment with their exquisitely motile processes. Intriguingly, microglia physically interact with neurons in normal and diseased brain. Recent studies showed that neuronal hyperactivity induced microglial interaction with neuronal somata and dendrites (*1–3*). In turn, microglia have also been observed to increase or decrease synaptic strength to maintain appropriate neuronal activity levels (*4–6*). The correlation between the reduced microglial dynamics and increased seizure activity in P2Y12 knockout mice suggest that microglia might be able to dampen the neuronal activity in the epileptic brain (*1*). Indeed, a recent study found that microglial contact of neuron downregulated its excitability in the zebrafish brain (*7*). However, it remains to be elucidated whether microglial modulation of neuronal hyperactivity is a general mechanism, and if so, what is the consequence of the spinal microglia-neuron communication in animal pain behaviors.

Although a wealth of evidence suggests that spinal microglia may play a critical role in chronic models of neuropathic pain within the time scale of days (*8*), involvement of microglia in acute peripheral inflammation-induced pain has been questioned (*9*). Microglial activation under neuropathic pain conditions has been demonstrated by altered morphology (*10*), increased expression of surface markers (such as CD11b, P2X4 receptors, etc) (*11*), phosphorylation of P38 MAP kinases and Src-family kinases Lyn (*8, 11–13*). However, under formalin acute inflammatory pain conditions, there were no obvious changes in morphology or cell surface marker expression within an hour after the insult. Consistently, no report has been able to pinpoint the specific roles of resting microglia in acute inflammatory pain hypersensitivity so far. The typical criteria for microglial activation do not take microglial dynamics and electrophysiological parameters into account. Particularly, microglial processes interact with neuronal components constantly within the time scale of minutes in the brain (*14, 15*). Therefore, whether microglia are functionally activated and contribute to inflammatory pain needs to be re-evaluated.

To address this question, we employed *in vivo* two-photon imaging to monitor microglial dynamics as well as whole-cell patch clamp recordings to detect microglial activation after intraplantar formalin injection. Moreover, using transgenic mice that enable us to specifically ablate resident microglia in a controllable fashion, we found that microglia are required for formalin-induced inflammatory pain. We further identified that microglial P2Y12 receptors participated in microglial dynamics, neuronal hyperactivity, and acute inflammatory pain. Our results demonstrate that increased microglial dynamics may amplify neuronal excitability underlying acute, sustained, inflammatory pain.

## RESULTS

### Ablation of CX_3_CR1^+^ cells diminished intraplantar formalin injection induced inflammatory pain

It is still unclear whether spinal microglia are engaged in acute peripheral inflammation-induced pain (*9*), since there are no obvious changes in microglial morphology within a few hours after the insult. CX_3_CR1 is predominantly expressed by microglia in the CNS, but is also found in a subset of monocytes, macrophages, natural killer cells, and dendritic cells in the periphery (*16*). To study the role of CX_3_CR1^+^ cells in acute inflammatory pain, we used a strategy to specifically express the diphtheria toxin receptor (DTR) in CX_3_CR1^+^ cells by crossing CX_3_CR1^CreER^ mice with Rosa26-stop-DTR mice (CX_3_CR1^CreER/+^:R26^iDTR/+^), and inducing expression with tamoxifen (TM) injections, as described previously (*17, 18*). TM (150 mg/kg in corn oil, 4 doses with 2 day intervals) or corn oil control was intraperitoneally (i.p.) injected to CX_3_CR1^CreER/+^:R26^iDTR/+^ mice and then two doses of diphtheria toxin (DT, i.p., 50 µg/kg, 2.5 µg/ml in PBS) were given at 3 and 5 days after last TM treatment to ablate CX_3_CR1^+^ cells, termed here as *CX_3_CR1^+^ Cell Ablation* (Fig. 1a). At 2 days after last DT injection, we examined the depletion efficiency of spinal cord microglia and dorsal root ganglia (DRG) macrophages by immunostaining for Iba1, as well as hind paw skin macrophages by immunostaining for F4/80 (Fig. 1a-b). In CX_3_CR1^CreER/+^: R26^iDTR/+^ mice without TM induced DTR expression (Control group), the microglia in the spinal cord dorsal horn and resident macrophages in DRGs and hind paw skins remain intact. However, in CX_3_CR1^CreER/+^: R26^iDTR/+^ mice with both TM and DT injection (CX_3_CR1^+^ Cell Ablation group), spinal microglia, DRG macrophages and hind paw skin macrophages were largely depleted (Fig. 1b-c).

**Figure 1.**
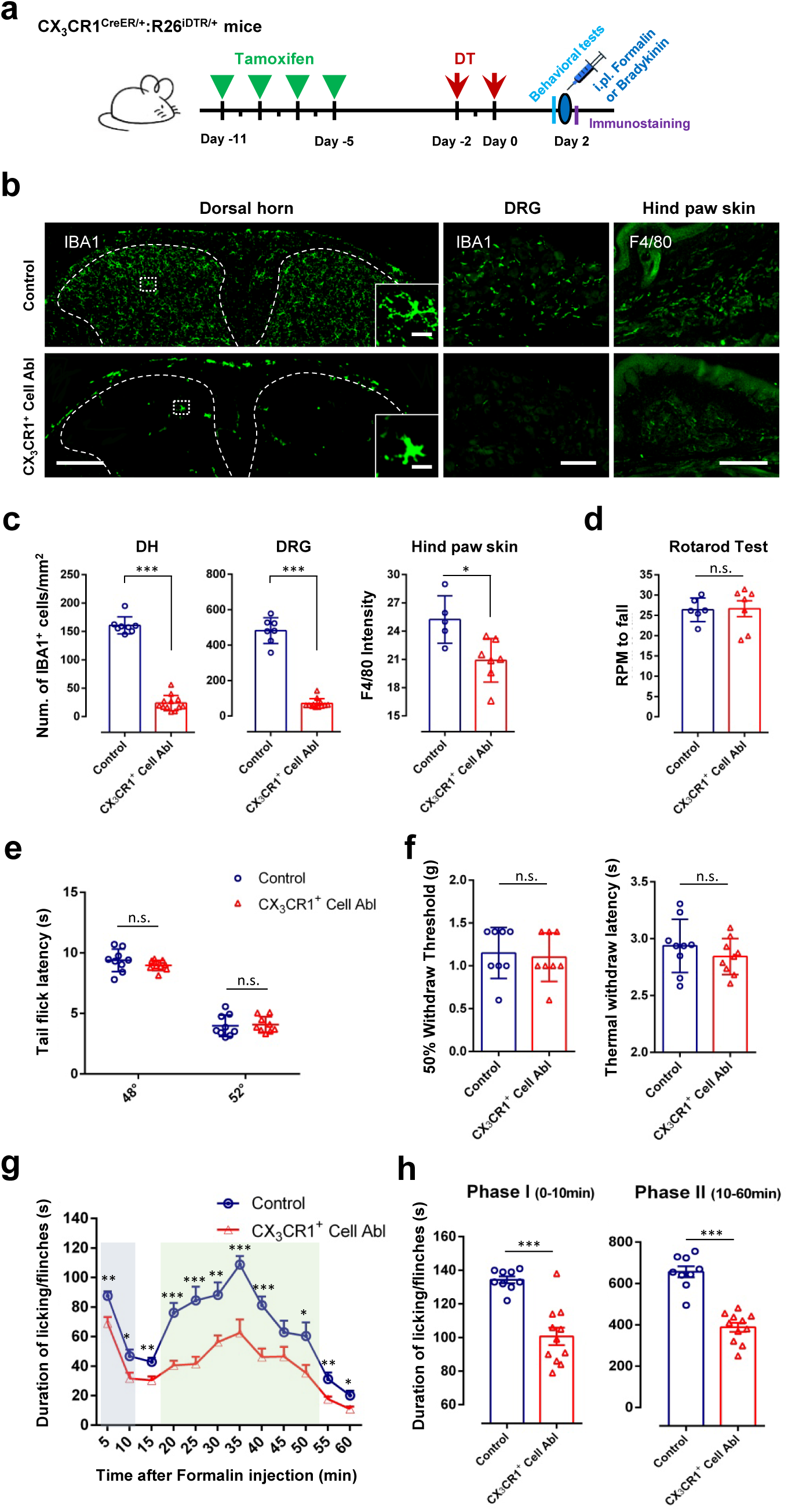
CX_3_CR1^+^ cell depletion reduces both formalin induced phase-I acute inflammatory pain and phase-II persistent inflammatory pain. **(a)** Schematic diagram showing the timeline of tamoxifen (TM) and diphtheria toxin (DT) administration, intraplantar injection of formalin (5% in PBS, 10 μl) or bradykinin (3 μg in PBS, 10 μl), behavioral tests (pain thresholds and rotarod test) and immunostaining in mice. **(b)** Iba1-positive cells were largely depleted 2 days after last DT injection in spinal cord dorsal horn (DH) and dorsal root ganglia (DRG). F4/80-positive macrophages in hind paw skin were also largely depleted at POD2 after the last DT injection, compared to the control group (mice without DT treatment). Left column, low magnification view of Iba1-positive cells in spinal cord DH of CX_3_CR1^CreER/ +^: R26^iDTR/+^ mice in control and CX_3_CR1^+^ cell ablation (Abl) mice. Inset: representative higher magnification images showing detailed microglia morphology. Scale bars represent 200 μm and 20 μm. Middle column, representative images of Iba1-positive cells in DRG in control and CX_3_CR1^+^ cell ablation mice. Scale bar, 100 μm. Right column, representative images of F4/80-positive macrophages in hind paw skin in control and CX_3_CR1^+^ cell ablation mice. Scale bar, 200 μm. **(c)** Quantitative data showing the density of Iba1-positive cells in the DH (***p < 0.001, unpaired 2-tailed Student’s t test, n = 8–12 mice/group), DRG (***p < 0.001, unpaired 2-tailed Student’s t test, n = 7–12 mice/group) and F4/80-positive cells in hind paw skin (*p < 0.05, unpaired 2-tailed Student’s t test, n = 5–7 mice/group) from control and CX_3_CR1^+^ cell ablation mice. Data are presented as mean ± SEM. **(d-f)** CX_3_CR1^+^ cell ablation does not affect acute pain responses and basal motor function. **(d)** Analysis of rotarod test in control and CX_3_CR1^+^ cell ablation mice showing that CX_3_CR1^+^ cell ablation did not affect basal motor function. (n = 6–7 mice/group). **(e-f)** Basal pain threshold for heat sensitivity (Tail immersion tests, Thermal Withdrawal Latency with Hargreaves Method) and mechanical sensitivity (Mechanical Withdrawal Threshold with von Frey tests) in control and CX_3_CR1^+^ cell ablation mice. Data are presented as mean ± SEM. n.s., no significance, unpaired 2-tailed Student’s t test, n = 8-9 mice/group. **(g-h)** Time course (0–60 min) of formalin-induced spontaneous pain behavior (licking/flinching) in control and CX_3_CR1^+^ cell ablation mice, as measured in every 5 min **(g).** Histogram representing formalin-induced Phase-I (1–10 min) and Phase-II inflammatory pain responses (10–60 min) in control and CX_3_CR1^+^ cell ablation mice. Both formalin-induced Phase-I and Phase-II inflammatory pain are reduced in CX_3_CR1^+^ cell depleted mice **(h)**. Data are presented as mean ± SEM; *P < 0.05, \*\**P* < 0.01, \*\*\**P* < 0.001, compared to control mice, unpaired 2-tailed Student’s t test, n = 9–11 mice/group.

Firstly, we examined the effect of CX_3_CR1^+^ cell ablation on basal motor and nociceptive responses in mice. We found that the mice devoid of CX_3_CR1^+^ cells show normal motor coordination in the rotarod test (Fig. 1d). In addition, there was no significant difference in basal pain threshold for heat sensitivity (such as tail immersion tests, thermal withdrawal latency tested with Hargreaves method) and mechanical sensitivity (such as mechanical withdrawal threshold tested with von Frey tests) between control and CX_3_CR1^+^ cell ablation mice (Fig. 1e-f).

Next, we examined the role of CX_3_CR1^+^ cells in formalin-induced pain, a well-established model of acute inflammatory pain. Formalin (5% in PBS, 10 μl) was injected into hind paw (intraplantar). In control mice, subcutaneous injection of formalin into the plantar side of a hind paw induced licking/flinch responses to the injected paw, typically concentrated in two distinct phases: acute phase I (0−10 min) and sustained phase II (10−60 min). We found that both phase I and phase II inflammatory pain behavioral responses were largely reduced in CX_3_CR1^+^ cell ablation mice (Fig. 1g). To investigate the sex-dependent role of spinal microglia in formalin-induced inflammatory pain, male and female mice were tested. We found that CX_3_CR1^+^ cell ablation significantly reduced formalin pain (both phase I and II) in either male and female mice (**Fig. S1a-b**). In addition to the formalin pain model, we further tested the effect of CX_3_CR1^+^ cell ablation in another inflammatory pain model induced by bradykinin. We found that the depletion of CX_3_CR1^+^ cells also attenuated pain behaviors induced by intraplantar bradykinin injection (3 μg in PBS, 10 μl) (**Fig. S2a**). Together, these results demonstrated that the ablation of CX_3_CR1^+^ cells reversed inflammatory pain behaviors.

### Deletion of peripheral macrophage/monocytes by clodronate reduces formalin-induced phase I acute inflammatory pain

Our above results demonstrate the pivotal function of CX_3_CR1^+^ cells in inflammatory pain. However, the respective role of peripheral monocytes/macrophages and central microglia in inflammatory pain hypersensitivity remains unknown. To address this question, clodronate liposomes were used to deplete peripheral tissue phagocytes including monocytes and macrophages (*19, 20*), termed here as *Clodronate Depletion*. Indeed, F4/80^+^ macrophages were largely depleted in hind paw skin 2 days after the last clodronate liposome treatment (Fig. 2a-b). In addition, F4/80^+^ macrophages in the spleen and liver were largely depleted after clodronate treatment (**Fig. S3**). However, Iba1^+^ microglia in spinal cord DH and Iba1^+^ macrophages in the DRGs were both preserved (Fig. 2a-b). Thus, clodronate liposomes were able to deplete hind paw skin macrophages but cannot ablate DRG macrophages or spinal microglia.

**Figure 2.**
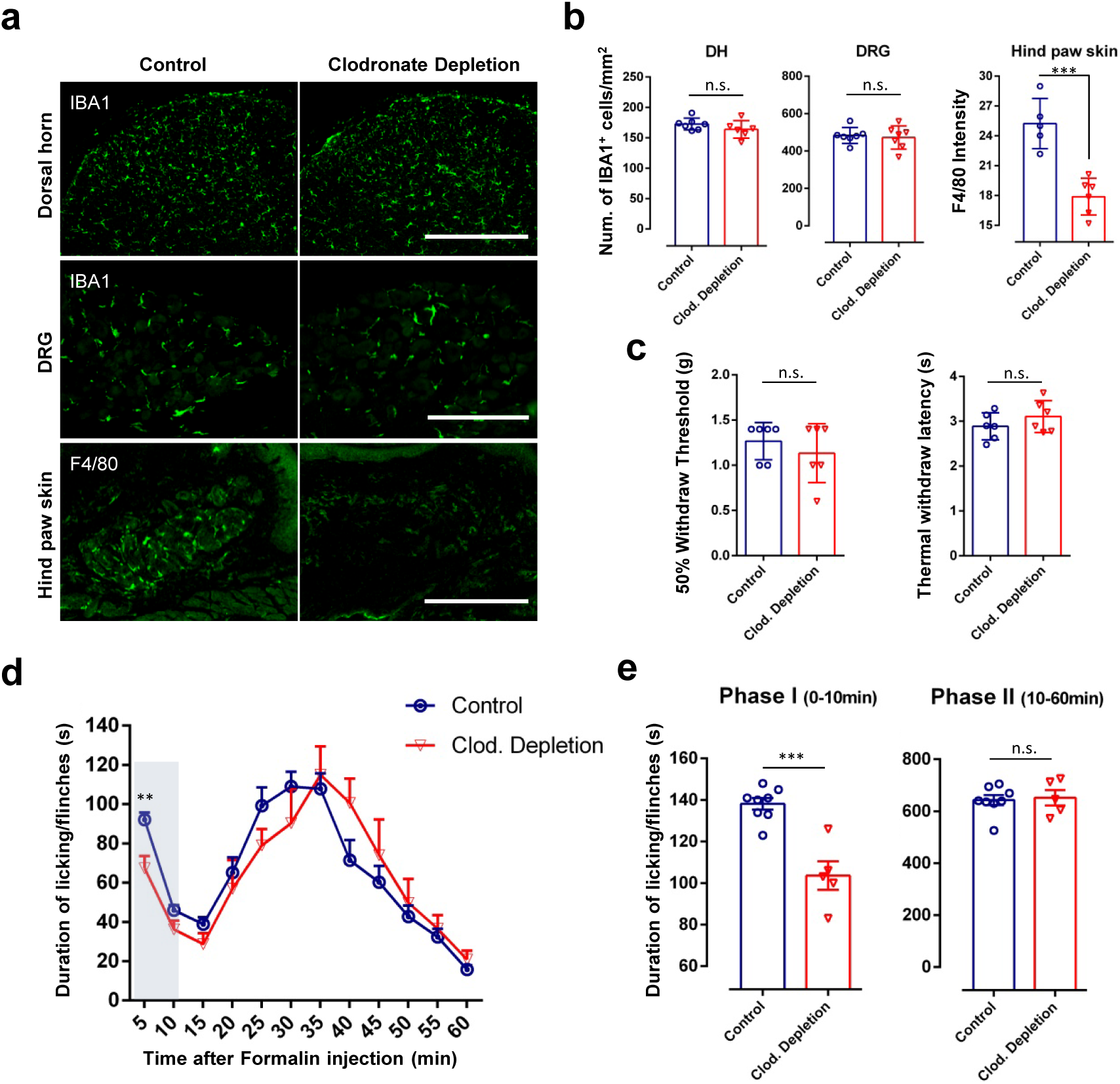
Depletion of peripheral macrophage reduces formalin induced phase-I acute inflammatory pain. **(a)** F4/80-positive cells were largely depleted in hind paw skin 2 days after clodronate liposome treatment (15 ml/kg, i.p.), while Iba1-positive cells in spinal cord DH and DRG were preserved. Representative images of Iba1-positive cells in DH and DRG and F4/80-positive macrophages in hind paw skin from the control and clodronate depletion mice. Scale bars represent 400 μm, 200 μm and 300 μm. **(b)** Quantitative data showing the density of Iba1-positive cells in DH, DRG (n.s., no significance, unpaired 2-tailed Student’s t test, n = 6–7 mice/group) and F4/80-positive cells in hind paw skin (***p < 0.001, unpaired 2-tailed Student’s t test, n = 5-6 mice/group) from control and clodronate (Clod.) depletion mice. Data are presented as mean ± SEM. **(c)** Basal pain threshold for heat sensitivity (Thermal Withdrawal Latency with Hargreaves Method) and mechanical sensitivity (Mechanical Withdrawal Threshold with von Frey tests) in control and clodronate depletion mice. n = 6 mice/group. Data are presented as mean ± SEM. **(d)** Time course (0–60 min) of formalin-induced spontaneous pain behavior (licking/flinching) in control and clodronate depletion mice, as measured every 5 min. Histogram representing formalin-induced Phase-I (1–10 min) and Phase-II (10–60 min) inflammatory pain responses in control and clodronate depletion mice. Note formalin-induced Phase-I acute inflammatory pain is reduced in clodronate depletion mice, while formalin-induced Phase-II inflammatory pain was not significantly affected. Data are presented as mean ± SEM, \*\**P* < 0.01, \*\*\**P* < 0.001, compared to control mice; unpaired 2-tailed Student’s t test, n = 5–8 mice/group.

We then examined the effect of clodronate liposome on pain behaviors. We found that depletion of peripheral tissue macrophages by clodronate did not affect basal nociceptive responses (Fig. 2c). However, formalin-induced phase I acute inflammatory pain behaviors were significantly attenuated following clodronate depletion (Fig. 2d). Interestingly, formalin-induced phase II inflammatory pain was not altered in clodronate-treated mice (Fig. 2d). Therefore, depletion of peripheral tissue macrophages inhibited acute phase I inflammatory pain induced by formalin. These results suggest that formalin-induced phase I pain is mainly augmented by skin macrophages important in the immediate inflammatory response.

### Ablation of resident microglia reduces formalin-induced phase-II persistent inflammatory pain

To further dissect the respective role of peripheral monocytes/macrophages and resident microglia in inflammatory pain, we again used CX_3_CR1^CreER/+^:R26^iDTR/+^ mice, which enabled us to exclusively deplete resident microglia in the CNS but not peripheral macrophage/monocytes. Since the resident microglia show much slower turnover (*21, 22*) while blood CX_3_CR1^+^ cells are replenished frequently, we were able to only ablate resident microglia by DT injection (50 µg/kg, 2.5 µg/ml in PBS) 3 weeks after TM application (i.p. 150 mg/kg) in CX_3_CR1^CreER/+^:R26^iDTR/+^ mice, termed here as *Microglia Ablation* (Fig. 3a). As expected, we found that spinal microglia were largely ablated 2-4 days after the last DT injection (Fig. 3b-c). Interestingly, we found that resident macrophages in DRGs can also be ablated acutely along with resident microglia in the spinal cord. However, DRG macrophages repopulated faster than spinal microglia. Three days after the last DT injection, the number of DRG macrophages reached similar levels as control animals while spinal microglia were still largely absent (Fig. 3b-c). In addition, macrophages in hind paw skin remained at similar levels (Fig. 3d). Macrophages in the liver and spleen and monocytes in the blood were also comparable between control and microglia ablation mice (**Fig. S4**). These results indicate that spinal resident microglia, but not peripheral macrophages can be selectively depleted using this approach.

**Figure 3.**
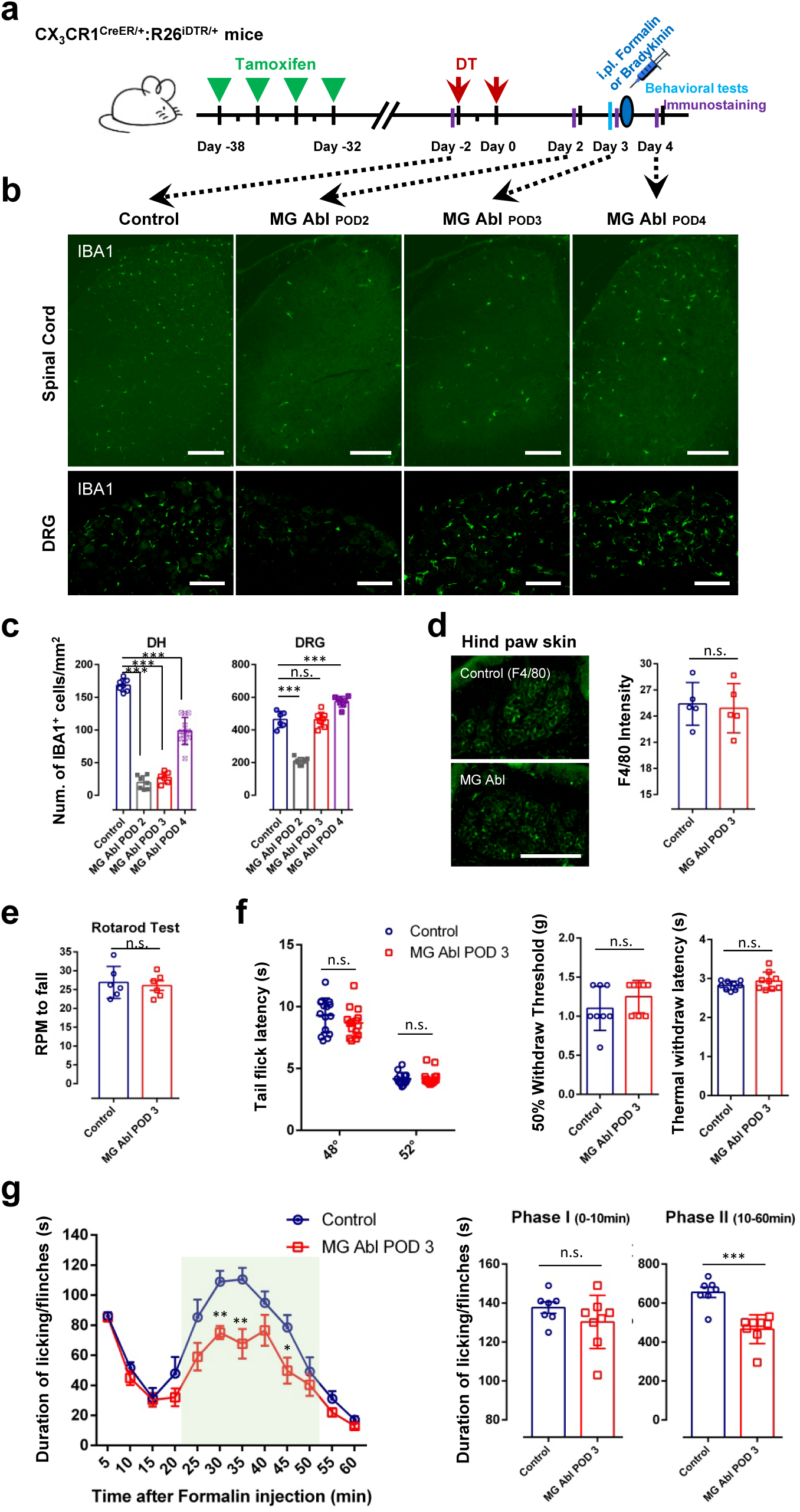
Ablation of resident microglia reduces formalin induced phase-II persistent inflammatory pain. **(a)** Schematic diagram showing the timeline of TM and DT administration, intraplantar injection of formalin (5% in PBS, 10 μl) or bradykinin (3 μg in PBS, 10 μl), behavioral tests (pain thresholds and rotarod test) and immunostaining in mice. Ablation or repopulation efficiency were checked at four time points before DT injection and days 2,3, 4 after the last DT injection. **(b-d)** Iba1-positive resident macrophages in DRGs can also be ablated acutely along with resident microglia in the spinal cord. However, DRG macrophages repopulated faster than that of spinal microglia. The number of DRG macrophages reached control levels 3 days after last DT injection, while spinal microglia were still largely absent. F4/80-positive macrophages in hind paw skin remain preserved after TM and DT treatment. **(b)** Representative images of Iba1-positive cells in spinal cord and DRG before DT injection and 2,3, 4 days after last DT injection. Scale bar, 200 μm. **(c)** Quantitative data showing the density of Iba1-positive cells in the DH (***p < 0.001, unpaired 2-tailed Student’s t test, n = 6–12 mice/group) and DRG (***p < 0.001, unpaired 2-tailed Student’s t test, n = 6–8 mice/group) from control and microglia ablation (MG Abl) mice. Data are presented as mean ± SEM. **(d)** Representative images of F4/80-positive macrophages in hind paw skin in control and microglia ablation POD3 mice. Scale bar, 200 μm. Quantitative data showing the density of F4/80-positive macrophages in skin from control and microglia ablation mice. (n.s., unpaired 2-tailed Student’s t test, n = 5 mice/group). **(e)** Analysis of rotarod test in control and microglia ablation mice indicating that CNS microglia ablation did not affect basal motor function (n = 6 mice/group). **(f)** Basal pain threshold for heat sensitivity (Tail immersion tests, n = 6 mice/group; Thermal Withdrawal Latency with Hargreaves Method, n = 9 mice/group) and mechanical sensitivity (Mechanical Withdrawal Threshold with von Frey tests, n = 8 mice/group) in control and microglia ablation POD3 mice. Data are presented as mean ± SEM; n.s., unpaired 2-tailed Student’s t test. **(g)** Time course (0–60 min) of formalin-induced spontaneous pain behavior (licking/flinching) in control and microglia ablation mice, as measured every 5 min. Histogram representing formalin induced Phase-I (1–10 min) and Phase-II inflammatory pain responses (10–60 min) in control and microglia ablation mice. Formalin-induced Phase-II persistent inflammatory pain is reduced in microglia ablation mice at POD3, while formalin-induced Phase-I acute inflammatory pain was not affected. Data are presented as mean ± SEM; *P < 0.05, \*\**P* < 0.01, \*\*\**P* < 0.001, compared to control mice; unpaired 2-tailed Student’s t test, n = 7–8 mice/group.

Next, we tested the impact of resident microglia ablation on pain behaviors. We found no significant difference in motor coordination following ablation or basal pain threshold (both for heat sensitivity and mechanical sensitivity) compared to controls (Fig. 3e-f). In the formalin-induced inflammatory pain model, there was no significant difference in phase I acute inflammatory pain between microglial ablation and control mice. However, behavioral responses during the late phase (phase-II persistent inflammatory pain behavioral responses) were significantly attenuated in microglial ablation mice (Fig. 3g). Consistent with the idea that microglia are critical for persistent inflammatory pain, we found that microglia ablation also reduced pain behaviors induced by intraplantar bradykinin injection (3 μg in PBS, 10 μl) (**Fig. S2b**). Thus, these results indicate a critical role for resident microglia in formalin-induced phase II persistent inflammatory pain or bradykinin-induced inflammatory pain.

### Increased dynamics of microglial processes during formalin induced phase-II persistent inflammatory pain

Microglia constantly survey the microenvironment in the CNS (*15, 23, 24*), but the dynamics of microglia in response to pain stimulation in vivo have not been well explored (*3*). To this end, we performed two-photon imaging of microglia in anesthetized CX_3_CR1^GFP/+^ mice, in which microglia were selectively labeled with GFP (*16*). Under control conditions, microglia have relatively stationary somata, but their processes dynamically sample the spinal cord parenchyma. After formalin injection (5% in PBS, 10 μl) in the hind paw, we found that microglial process motility was significantly enhanced starting 10 min after the insult. The increased motility index lasted for at least 30 minutes (Fig. 4a-c). In addition, the total process length of microglia gradually increased after formalin injection (Fig. 4d). Consistently, the complexity of microglia was also enhanced alongside the elongation, as shown by Sholl analysis (Fig. 4e). To further confirm that the increased microglial motility was not due to the *in vivo* imaging procedures or anesthetics, we examined microglial morphology in fixed spinal cord slices before and 1 h after formalin injection. Indeed, we found that the number of primary processes and microglia cell area was increased after formalin injection (**Fig. S5**). Taken together, our results demonstrate that microglial process dynamics were increased during acute, sustained inflammatory pain induced by formalin injection.

**Figure 4.**
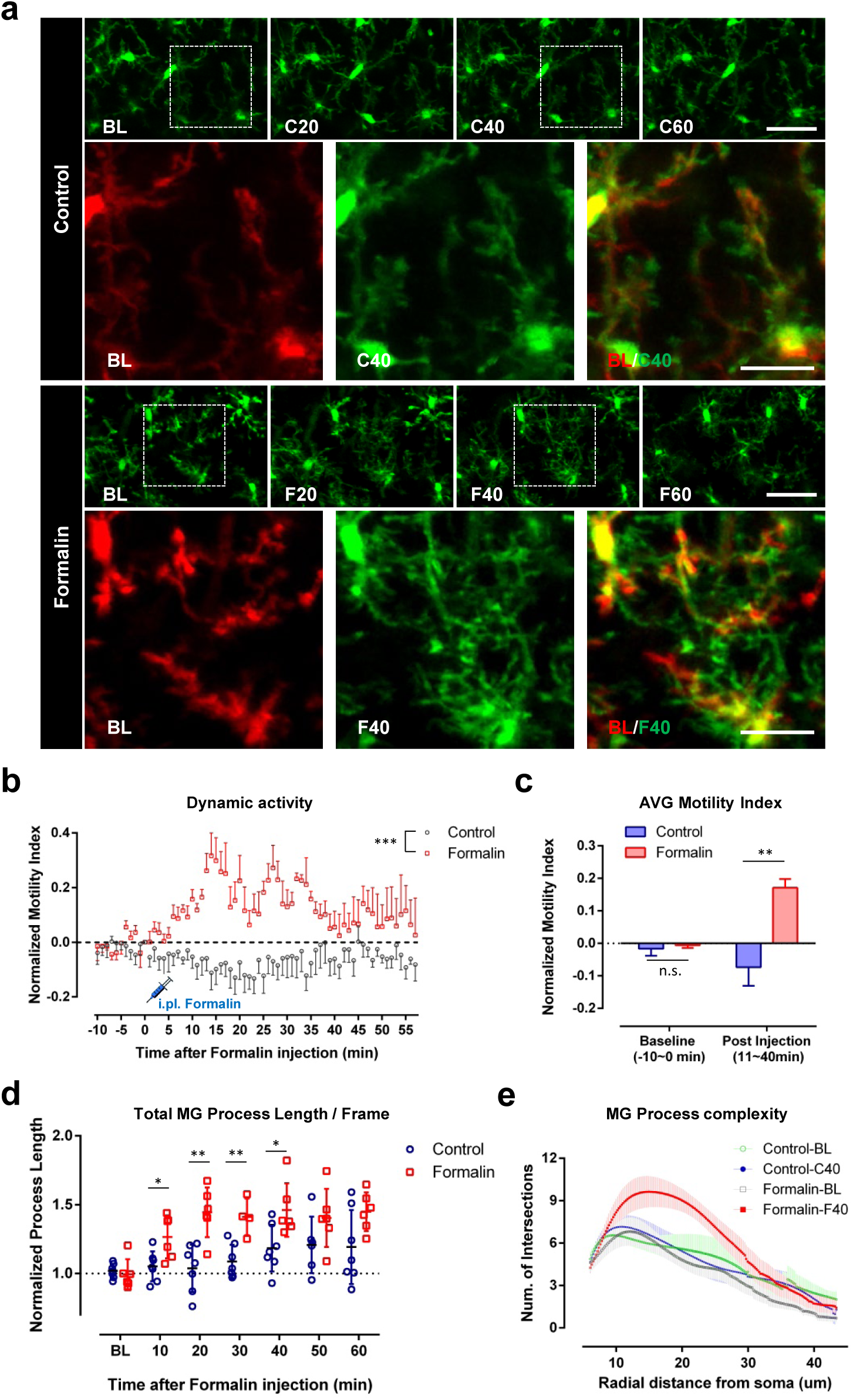
Microglial process dynamics are enhanced during formalin induced phase-II persistent inflammatory pain. **(a)** Representative *in vivo* z-stack images of L4-5 ipsilateral spinal microglia from time-lapse imaging taken before (–10 min, BL) and 20 (F20), 40 (F40), or 60 min (F60) after either intraplantar formalin injection (5% in PBS, 10 μl) or control injection (PBS). Higher magnification images from boxed regions of BL in red and F40 in green were merged in BL/F40 to show microglial process extension. Time lapse two-photon z-stack images were also taken from control mice at before (–10 min, BL) and 20 min (C20), 40 min (C40), 60 min (C60) after intraplantar saline injection. Higher magnification images from boxed regions of BL in red and C40 in green were merged in BL/C40. Note that there was no dramatic change in microglial process extension in control conditions. Scale bars represent 40 μm and 20 μm. **(b)** Quantitative data showing dynamics of spinal microglial processes, represented as normalized motility index, in control and formalin-treated groups. Data are shown as mean ± SEM. ***P < 0.001, compared to control. Two-way ANOVA. n = 5-6 mice/group (Control: n = 6, Formalin: n = 5). **(c)** Quantitative data showing average motility index of spinal microglial processes in control and formalin-treated group within different time windows (−10 ∼ 0 min before and 11 ∼ 40 min after saline/formalin injection). Data are shown as mean ± SEM. **P < 0.01, compared to control. Unpaired 2-tailed Student’s t test. n = 6 for control and n = 5 for formalin-treated group. **(d)** Quantitative data showing spinal microglial process length, represented as normalized process length, in control and formalin-treated groups at different time points with respect to saline/formalin injection. Process lengths were analyzed every 10 min. Data are shown as mean ± SEM. *P < 0.05, **P < 0.01, compared to control. Unpaired 2-tailed Student’s t test. n = 6-7/group. **(e)** Sholl analysis plot representing microglia complexity in control and formalin-treated group. n = 7-13/group.

Formalin is known to activate TRPA1 channels in nociceptors causing their hyperactivity (*25*). To directly test whether increased peripheral neuronal activity is sufficient to induce an increase in microglial dynamics, we employed high frequency stimulation in the sciatic nerve (4 trains of 100Hz stimulation (0.5 ms rectangular pulses at 20V) for a 2-sec duration, with a 10-sec inter-train interval) and monitored microglia motility (**Fig. S6**). We indeed found that HFS itself triggers increased process motility in spinal microglia (**Fig. S6a-c**). Consistently, we also observed that HFS can increase process length and branch complexity in spinal microglia (**Fig. S6d-e**). Therefore, rapid stimulation of the sciatic nerve can mimic multiple aspects of formalin injection in enhancing microglial process dynamics in the spinal cord.

### Electrophysiological activation of spinal microglia after formalin-induced inflammatory pain

The increased microglial dynamics in response to formalin pain stimulation is surprising, as reactive microglia were only seen 2-3 d after formalin injection and 1 d after peripheral nerve injury (*13, 26*). To directly test microglial reactivity, we also performed whole-cell patch clamp recordings in spinal microglia 40 min after intraplantar formalin injection. Under normal conditions, spinal microglia show minimal current responses to membrane depolarization (Fig. 5a), which is consistent with quiescent microglia in the brain (*27*). However, spinal microglia from formalin-injected mice show significantly larger outward-rectifying K^+^ currents in response to membrane depolarization (Fig. 5b). We observed similar currents in activated spinal microglia 3 d after peripheral nerve injury (*3*). In addition, it is also known that activated microglia after facial nerve axotomy bear similar outward-rectifying K^+^ currents (*28*). These results indicate that spinal microglia may display electrophysiological hallmarks of activation well before any morphological changes can be observed following formalin injection.

**Figure 5.**
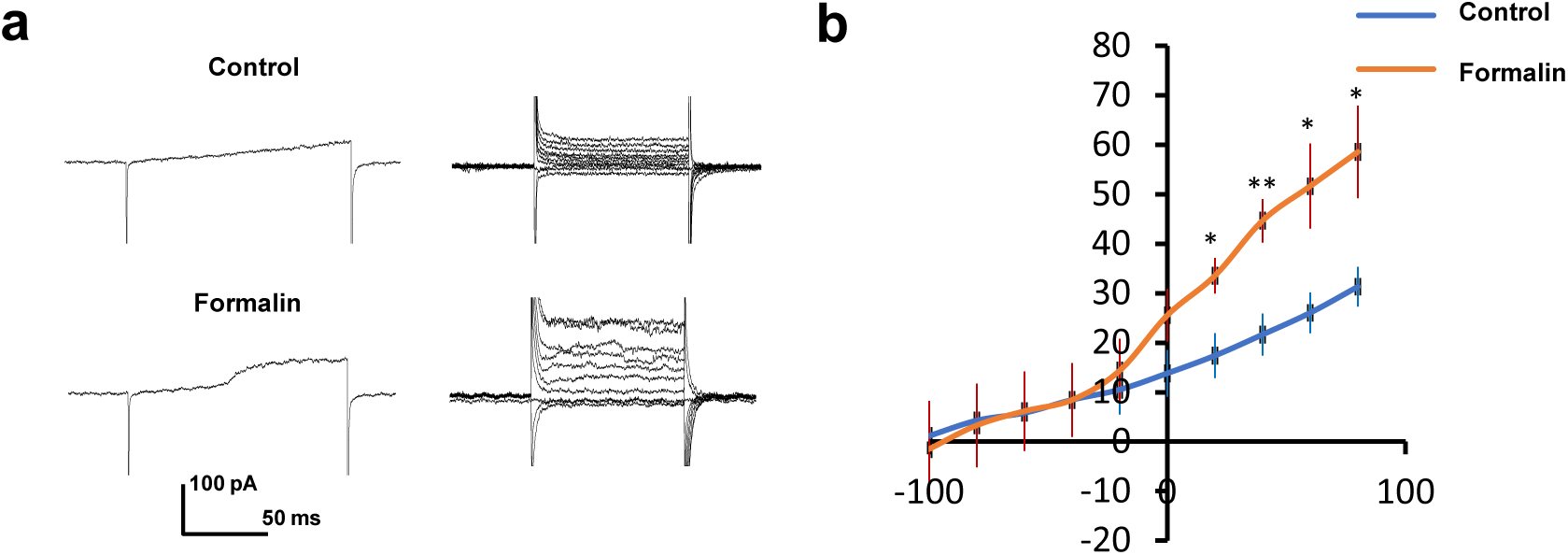
Electrophysiological properties of microglia during formalin induced phase-II persistent inflammatory pain. **(a)** Representative traces showing whole cell recording of microglial currents in response to ramp from −100 mV ∼ +80 mV (left) or depolarization steps from −100 mV to 80 mV (right) in control and formalin-treated groups. **(b)** Quantified summaries of microglial current in response to depolarization steps from −100 mV to 80 mV from control and formalin-treated mice. Holding potential: −60 mV. n = 5 cells/group. Data are shown as mean ± SEM. *P < 0.05; **P < 0.01, compared to control.

### Microglia ablation reduces neuronal hyperactivity in the spinal cord after formalin-induced inflammatory pain

Microglial dynamics and electrophysiological activation may affect neuronal activity during formalin-induced inflammatory pain. To test this idea, we examined the expression of activity-dependent immediate early genes c-fos and phosphorylated ERK (pERK). Intraplantar injection of formalin induced abundant c-fos and pERK expression co-localized with NeuN immunostaining (Fig. 6a-b), suggesting that formalin induced neuronal hyperactivity in the spinal dorsal horn. Next, we compared the formalin-induced c-fos and pERK expression in control and microglia ablation mice. We found that microglia ablation significantly decreased c-fos expression in spinal dorsal horn following intraplantar formalin injection (Fig. 6c-d). Consistent with our c-fos results, microglia ablation also reduced pERK levels following formalin injection (Fig. 6e-f). Therefore, neuronal activity in the spinal cord triggered by formalin injection was attenuated in the absence of microglia. Together, these results suggest that microglia might be able to amplify neuronal activity in the spinal cord during formalin-induced inflammatory pain.

**Figure 6.**
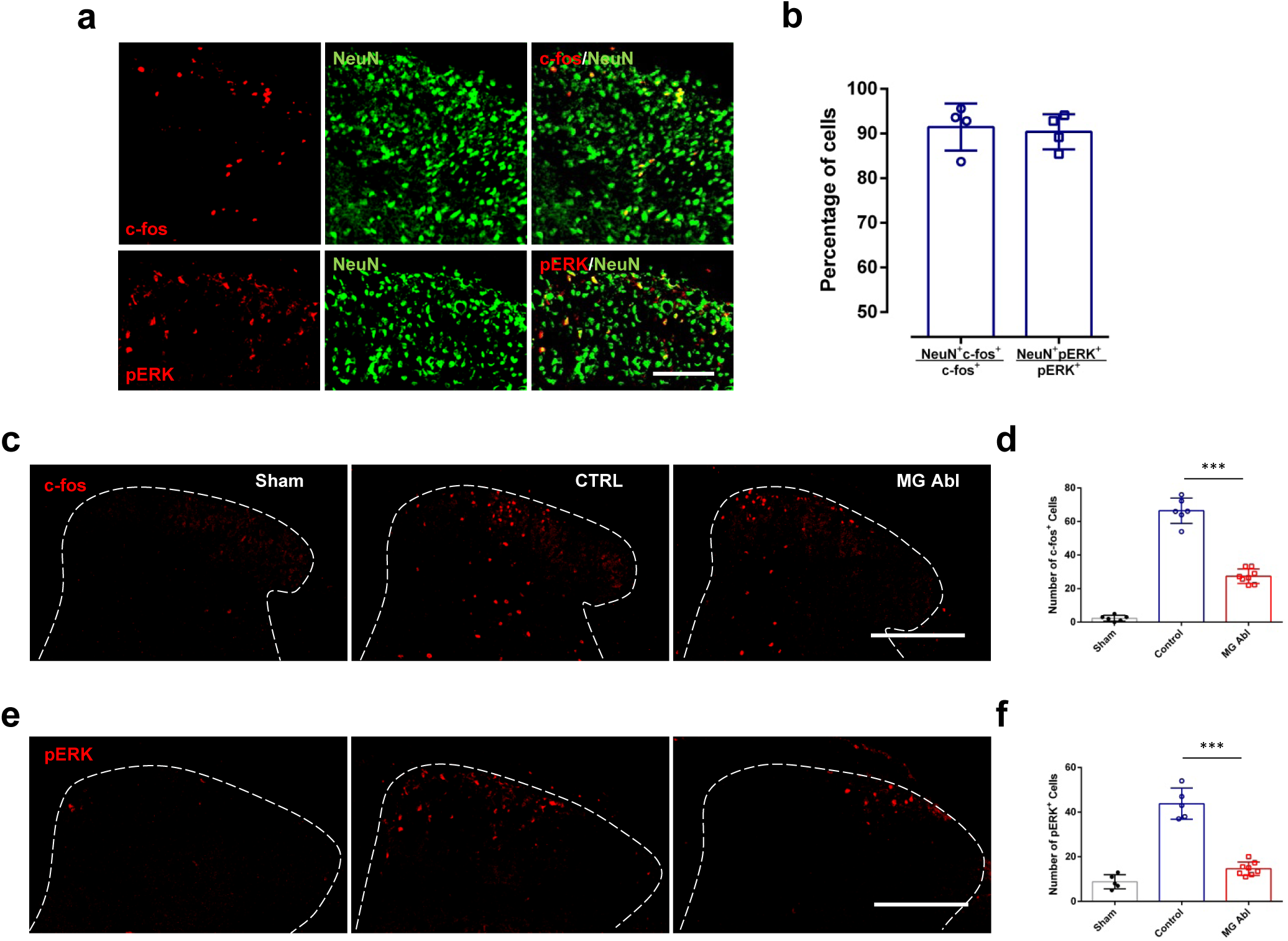
Reduced formalin-induced neuronal activity in the absence of microglia in the spinal cord. **(a-b)** Intraplantar injection of formalin induced c-fos expression and pERK activation co-localized with NeuN immunostaining. **(a)** Representative images showing double staining of c-fos^+^ or pERK^+^ (red) and NeuN^+^ (green) cells in the spinal cord DH either 2 hours after formalin injection (c-fos staining) or 40 min after formalin injection (pERK staining). Scale bar, 100 mm. **(b)** Quantification of the percentage of c-fos^+^NeuN^+^ or pERK^+^NeuN^+^ cells among among all c-fos^+^ or pERK^+^ cells. n = 4 mice/group. **(c, e)** Representative images of c-fos -positive **(c)** and pERK -positive **(e)** cells in ipsilateral dorsal horn from the sham, control (CTRL) and microglia ablation (MG Abl) POD3 mice. Scale bar, 200 μm. **(d, f)** Quantitative data showing the number of c-fos^+^ cells 2 hours after intraplantar formalin injection **(d)** or pERK^+^ cells 40 min after intraplantar formalin injection **(f)** in the ipsilateral dorsal horn of WT sham, WT control and microglia ablation POD3 mice. Data are presented as mean ± SEM; ***P < 0.001, compared with control, unpaired 2-tailed Student’s t test, n = 5–8 mice/group.

### P2Y12 receptor in microglia dynamics, neuronal excitability and formalin inflammatory pain

The P2Y12 receptor is specifically expressed in microglia (*3, 29*) and it controls microglial interactions with hyperactive neurons under seizure conditions (*1*). Here we wanted to investigate whether microglial P2Y12 participates in the increased process dynamics induced by intraplantar formalin. Acute spinal cord slices were used to image microglial dynamics. Consistent with our *in vivo* results, microglia exhibited increased motility 40 min after intraplantar formalin injection in wild-type mice (Fig 7a-b). However, formalin-induced increases in microglial process dynamics were abolished in P2Y12^-/-^ slice (Fig 7a-b). These results demonstrate that altered microglial dynamics following formalin injection are dependent on microglial P2Y12 signaling.

**Figure 7.**
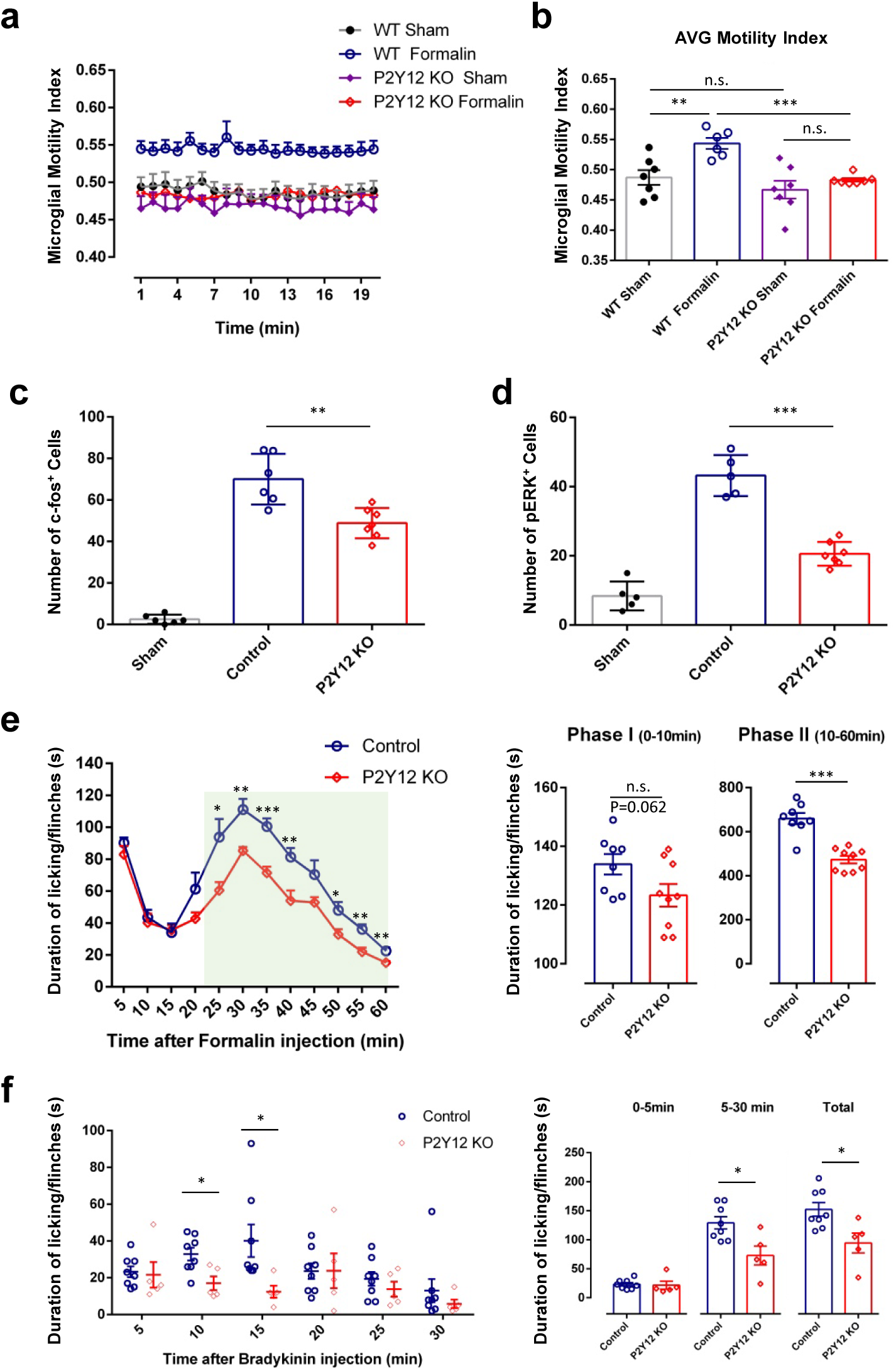
P2Y12 receptor in microglia dynamics, neuronal excitability and formalin induced phase-II persistent inflammatory pain. **(a-b)** Quantification of microglial motility indices at 40 min after intraplantar injection from WT and P2Y12 knockout (KO) spinal cord slices over a 20 min imaging session **(a)** or as the group average for each condition **(b)**. Sham, intraplantar saline injection; Formalin, intraplantar formalin injection. Data are shown as mean ± SEM. **P < 0.01, ***P < 0.001, unpaired 2-tailed Student’s t test, n = 6-7 slices from 3 to 5 mice per group for slices images. **(c-d)** The number of c-fos^+^ cells and pERK^+^ cells were largely reduced in P2Y12 KO mice compared with wild-type mice after formalin injection in the ipsilateral dorsal horn from the WT sham, WT control and P2Y12 KO groups. Data are presented as mean ± SEM; **P < 0.01, ***P < 0.001, compared with WT Control, unpaired 2-tailed Student’s t test, n = 5–7 mice/group. **(e)** Time course (0–60 min) of formalin-induced spontaneous pain behavior (licking/flinching) in WT control and P2Y12 KO mice, as measured every 5 min. Histogram representing formalin-induced Phase-I (1–10 min) and Phase-II inflammatory pain responses (10–60 min) in WT control and P2Y12 KO mice. Note the formalin-induced Phase-II inflammatory pain is significantly reduced in P2Y12 KO mice. Data are presented as mean ± SEM; *P < 0.05, **P < 0.01, ***P < 0.001, compared to WT control mice, unpaired 2-tailed Student’s t test, n = 8–9 mice/group. **(f)** Time course (0–30 min) of bradykinin-induced spontaneous pain behavior (licking/flinching) in WT control and P2Y12 KO mice, as measured every 5 min. Histogram representing bradykinin-induced inflammatory pain responses within 0–5 min, 5–30 min and 0-30 min in WT control and P2Y12 KO mice. Bradykinin-induced inflammatory pain is reduced within 5–30 min and 0-30 min in P2Y12 KO mice. Data are presented as mean ± SEM; *P < 0.05, compared to WT control mice, unpaired 2-tailed Student’s t test, n = 5–8 mice/group.

To further address the functional significance of P2Y12-dependent process dynamics, we compared formalin-induced c-fos or pERK expression between wild-type and P2Y12^-/-^ mice. We found that the number of neurons expressing c-fos induced by formalin injection was largely reduced in P2Y12^-/-^ mice compared with wild-type mice (Fig 7c). In line with these results, there were fewer pERK^+^ cells in P2Y12^-/-^ mice than those in wild-type mice after intraplantar formalin injection (Fig 7d). Next, we compared the formalin inflammatory pain behaviors between wild-type and P2Y12^-/-^ mice. If P2Y12 controls microglia motility and subsequent neuronal excitability, we expected the pain behaviors to be attenuated in P2Y12^-/-^ mice. Indeed, formalin-induced inflammatory pain was significantly reduced, particularly during phase II observations in P2Y12^-/-^ mice (Fig. 7e). In addition, we examined bradykinin-induced inflammatory pain in P2Y12^-/-^ mice and found that P2Y12 deficiency reduced pain behaviors induced by bradykinin (3 μg in PBS, 10 μl) (Fig. 7f). Together, these results indicate that microglial P2Y12 receptors increase process dynamics, aggravate neuronal activity, and amplify inflammatory pain behaviors after intraplantar formalin or bradykinin injection.

## DISCUSSION

It is well established that spinal microglia participate in chronic neuropathic pain (*30, 31*). However, the role of microglia in acute inflammatory pain has not been carefully studied because microglial activation is debatable under these conditions (*32, 33*). Here we revisited this idea using *in vivo* imaging and whole-cell recordings of microglia in spinal cord after acute inflammatory pain. Additionally we used genetic mouse lines to selectively deplete resident microglia in a controllable fashion. We found that formalin injection increased microglia process dynamics and electrophysiological signatures of activation in the spinal dorsal horn. Depletion of resident microglia significantly dampened neuronal activity markers and phase-II persistent inflammatory pain induced by formalin injection. Finally, we determined that the microglial P2Y12 receptor is responsible for increased microglia dynamics, neuronal excitability, and formalin-induced pain behaviors. These novel findings demonstrated that spinal microglia are able to amplify neuronal activity and thereby participate in sustained inflammatory pain signaling.

### The role of spinal microglia in acute inflammatory pain

Activation of spinal microglia has been reported in several models of inflammatory pain. For instance, there is an increase in microglia activation markers in the lumbar spinal cord following injection of Complete Freund’s Adjuvant (CFA) into the rat hind paw (*34, 35*). Similarly, robust activation was observed in the dorsal horn ipsilateral microglia after formalin injection into the rat hind paw starting at 1-3 d and peaking at 7 d post injection (*35*). The expression of cyclooxygenase1 by microglia was observed from 1d −2w after formalin injection (*36*). Microglial activation can be defined in several ways, including, morphological change from ramified to amoeboid (*37*), increase in microglial numbers (*38, 39*) or an enhanced expression of microglial markers (*34, 40*). These signs of microglial activation usually appear 2-3 d after the peripheral inflammation or nerve injury and hence it was believed that microglial activation does not occur during acute phase of inflammatory pain (*32, 33*). In the current study, we detected significant changes in microglial dynamics and electrophysiological parameters, suggesting microglial activation occurs within 1-2 h following formalin injection. Consistent with our report, it was shown that peripheral formalin injection could phosphorylate microglia p38 within minutes without any distinct changes in microglial morphology (*41, 42*). Whether p38 phosphorylation links to increased microglial process dynamics or electrophysiological activation warrants further studies.

It has been previously shown that the formalin-induced phase I pain (occurring immediately post-injection) is caused by the direct stimulation of nociceptors, while the phase II pain (occurred after a short period of quiescence) reflected central sensitization (*43*). In order to dissect the role of CNS microglia and peripheral macrophages/monocytes in the formalin-induced acute inflammatory pain, we adopted an ablation strategy (*17, 18*). When CX_3_CR1 cells (including microglia, macrophages and monocytes) were ablated, we found that both phase I and II inflammatory pain were significantly attenuated. Using clodronate to deplete skin macrophages, we found that phase I acute inflammatory pain were reduced. More importantly, when microglia were specifically ablated by taking advantage of the difference in turnover rate of microglia and macrophages/monocytes (*17, 21*), we found that pain behavioral responses during phase II but not phase I following formalin injection were significantly diminished. Taken together, our results demonstrate that while skin macrophages seem involve in phase I acute inflammatory pain, resident microglia are specially for central sensitization critical for formalin-induced phase II persistent inflammatory pain.

In addition to the spinal microglia, supraspinal microglia are reported to be an important player in inflammatory pain (*44*). For example, microinjection of the glial inhibitors such as minocycline into the rostral ventromedial medulla (RVM) produced a time-related attenuation of behavioral hypersensitivity by hind paw inflammation (*45*). Since microglia were depleted in the entire spinal cord and brain in our ablation model, the involvement of supraspinal microglia in inflammatory pain has to be considered while interpreting the results. In addition, it should be taken into account that non-physiological death of microglia may by themselves alter the neuronal microenvironment causing a change in neuronal activity and therefore pain phenotype (*46*). In both CX3CR1 cell ablation and microglia ablation mice, we did not notice the obvious motor dysfunction in the current study or previous studies (*17, 18*), unlike a recent report suggesting the microglial ablation induce neurodegeneration and ataxia (*47*). The discrepancy could be due to the difference dose of DT used (high dose may cause unwanted side effects) or behavioral tests at different time points after DT ablation. Nevertheless, microglia depletion approaches are powerful to temporally and specifically dissect the role of resident microglia and peripheral macrophages in chronic pain.

### Microglial regulation of neuronal activity in the brain and spinal cord

Accumulating evidence suggests that microglia regulate neuronal activity in the brain (*1, 4, 5, 7*). Here, we first demonstrate that formalin-induced inflammatory pain is associated with the increase of microglial process dynamics as well as microglial K^+^ channel activation in the spinal dorsal horn. More importantly, we found that microglia seem to amplify neuronal signals by increase c-fos and pERK activity, and thereby facilitating persistent inflammatory pain behaviors. In zebrafish brain, microglia-contacted neurons displayed less activity than non-contacted neurons, suggesting that microglia has an inhibitory effect (*7*). Consistent with the idea, microglia may dampen neuronal hyperactivity under seizure context (*1, 4*) or even under physiological conditions (*48*). On the contrary, a recent study reported that microglial contact increased Ca^2+^ activity in the dendritic spines in mouse cortex (*5*). In addition, selective microglial activation in the spinal cord was sufficient to facilitate synaptic strength between primary afferent C-fibers and lamina I neurons (*49*). More recently, our results demonstrate that microglia promoted maladaptive synaptic plasticity in the spinal dorsal horn while impaired hippocampal synaptic plasticity after peripheral nerve injury (*50, 51*). Hence, microglia are thought to exert differential regulation of neuronal activity under various contexts.

The potential mechanisms of microglial regulation of neuronal activity may involve direct physical interaction or through the secretion of cytokine/chemokines such as CX_3_CL1, IL-1β, TNF-α, BDNF, etc. For example, microglial secretion of TNF-α was shown to regulate synaptic plasticity and thereby inflammatory pain following formalin/bradykinin injections (*52*). Similarly, peripheral nerve injury alters synaptic connectivity following mediated by microglia TNF-α/TNFR1 signaling (*50*). In addition, recently studies showed that activation of CX_3_CR1 receptor by fractalkine induced the release of IL-1β from microglia resulting in enhanced nociceptive pathways (*49*). Here, we found that spinal microglia are able to amplify neuronal activity, and microglia ablation reduced formalin-induced c-fos and p-ERK expression in spinal dorsal horn neurons, in a mouse model of acute inflammatory pain. Future studies are needed to understand the exact molecular mediators from microglia that regulate neuronal activity in acute inflammatory pain.

### Microglial purinergic signaling in acute inflammatory pain

Microglia-neuron interaction is bidirectional (*6, 46*) and the increased neuronal activity following inflammatory pain can alter the activation status of microglia. Indeed, we observed the increased microglial dynamics and electrophysiological activation after peripheral formalin injection. Here, we investigated the microglial purinergic pathway based on the critical role of P2Y12 receptor in microglia-neuron interactions (*1, 48, 53, 54*). We found that formalin-induced an increase in microglial process dynamics is inhibited in P2Y12^-/-^ mice indicative of reduced neuronal activity. Consistent with our findings, another study showed that activation of P2Y12 receptors in satellite glial cells is involved in the enhancement in neuronal activity in the trigeminal nucleus and nocifensive reflex behavior following lingual nerve injury (*55*). It worth noting that microglial P2Y12 receptor seems playing opposite function to neuronal activity in the brain and the spinal cord. P2Y12 deficiency in the brain increases neuronal activity that might be important for innate fear behaviors or seizures (*1, 48*). The differential functions of brain and spinal microglia may reflect microglial heterogeneity in different CNS regions (*50, 56*). Nevertheless, our study suggests that spinal microglia amplify neuronal activity in acute inflammatory pain; reciprocally, neuronal hyperactivity regulates microglial dynamics via the P2Y12 receptor.

In terms of pain phenotype, we found that formalin-induced persistent inflammatory pain was suppressed in P2Y12^-/-^ mice, similar to those microglia-depleted mice. Consistent with our results, it has been shown that intrathecal administration of P2Y12 receptor antagonists alleviated mechanical and thermal hyperalgesia following intraplantar CFA injection (*57*). In peripheral nerve injury model, P2Y12^-/-^ mice displayed impaired tactile allodynia without any change in basal mechanical sensitivity following L5 spinal nerve transection (*3, 58*). In addition, a single intrathecal injection of P2Y12 antagonists reversed the nerve-injury induced tactile allodynia (*58, 59*). Since P2Y12 receptor is also present in the platelets (*60*) and DRG satellite cells, future study is needed to use conditional knockouts of P2Y12 receptors to test the function of microglial specific P2Y12 receptor in inflammatory and neuropathic pain.

In conclusion, our results demonstrate that microglia play a key role in the formalin-induced phase-II persistent inflammatory pain by amplifying neuronal activity. This study has established that physiological microglia are not passive bystanders, but actively contribute to the central sensitization underlying acute sustained inflammatory pain. The involvement of P2Y12 receptor signaling in neuron-microglia interaction allows for the development of targeted therapies for ameliorating sustained inflammatory pain.

## MATERIALS AND METHODS

### Animals

Both male and female animals were used in this study in accordance with institutional guidelines, as approved by the animal care and use committee at Rutgers University, Fourth Military Medical University, and Mayo Clinic. All animals were housed under controlled temperature, humidity, and lighting (light - dark 12:12 h cycle) with food and water available *ad libitum*.

C57BL/6J mice, CX_3_CR1-GFP reporter mice and Rosa26-stop-DTR (R26^iDTR/+^) mice were purchased from the Jackson Laboratory. Age-matched C57BL/6J and Heterozygous GFP reporter mice (CX_3_CR1^GFP/+^) were used as a wild-type control. 7∼12 week old adult mice were used for inflammatory pain behavior observation, and spinal cord *in vivo* and slice 2-photon experiments. For the production of CX_3_CR1^CreER-EYFP/+^ mice, an original breeding stock was gifted by Dr. Wen-Biao Gan at New York University. The CX_3_CR1^CreER-EYFP/+^ mice were crossed with R26^iDTR/+^ mice to obtain CX_3_CR1^CreER/+^: R26^iDTR/+^ mice. For the production of P2Y12 knockout mice (P2Y12^-/-^), an original breeding stock was gifted by Dr. Michael Dailey at the University of Iowa. P2Y12^-/-^ mice were viable and showed no developmental defects. CX_3_CR1-GFP reporter mice were crossed with P2Y12^-/-^ mice to generate the P2Y12^-/-^;CX_3_CR1 ^GFP /+^ mice.

### CX_3_CR1^+^ cell ablation and microglia ablation

Tamoxifen (Cat. T5648 TM, Sigma) was given as a solution in corn oil (Cat. C8267 Sigma). Adult mice (over P42) received four doses of TM (150 mg/kg, 20 mg/ml in corn oil) at 48 hour-intervals by intraperitoneal injection (i.p.). For total CX_3_CR1^+^ cell ablation (*CX_3_CR1^+^ cell ablation*), two doses of Diphtheria Toxin (DT, Sigma, Cat. #D0564, 50 µg/kg, 2.5 µg/ml in PBS) was given at 3 and 5 days after the last TM treatment. For CNS CX_3_CR1^+^ ablation (*Microglia Ablation*), the interval between the last TM and the first DT was more than 3 weeks. Mice administered with DT only (without TM) were used as control for all CX_3_CR1^+^ cell ablation experiments.

### Clodronate depletion

In some experiments, the circulating/tissue monocyte/macrophage population was depleted by Liposome-encapsulated clodronate (*ClodronateLiposomes.org*), as described previously (*18*). Clodronate liposomes (15 ml/kg) were i.p. injected 2 d before formalin or bradykinin treatment.

### Formalin or bradykinin-induced hind paw acute inflammation pain

To produce acute inflammatory pain, formalin (Cat. F9037, sigma; 5% in PBS, 10 μl) or bradykinin (Cat. B3259, sigma; 3 ug in PBS, 10 μl) was injected into hind paw (intraplantar). PBS was used in sham-operated mice. Animals were habituated to the testing environment for at least 2 d before the testing. The experimenter scoring pain behaviors was blinded to treatment. We assessed formalin- and bradykinin-evoked spontaneous nociceptive reactions by measuring the time mice spent licking the affected paw or flinching every 5 min for 60 min the formalin condition, or every 5 min for 30 min in the bradykinin condition.

### Behavioral testing

#### Tail immersion withdrawal latency

We used the tail immersion-withdrawal test to determine the thermal pain/withdrawal threshold for lightly restrained mice (*61*). The distal third of the tail was immersed directly in water at a set temperature of either 48°C or 52°C. The thermal withdrawal latency was recorded to the nearest 0.01 s as a characteristicly vigorous tail reflex response. A maximum latency of 20 and 10 s was permitted at temperatures of 48°C and 52°C, respectively, to prevent tissue damage.

#### Hind paw thermal withdrawal latency

The basal pain threshold for heat sensitivity was assessed by measuring hind paw withdrawal latency. The heat stimulation was provided following a previously described protocol (*62*). An analgesia meter (IITC Inc life science) was used to provide a heat source. In brief, each mouse was placed in a box containing a smooth, temperature-controlled glass floor (30°C) and allowed to habituate for 20 min. The heat source was then focused on a portion of the hind paw, which was flush against the glass, and a radiant thermal stimulus was delivered to that site. The stimulus shut off when the hind paw moved (or after 6 s to prevent tissue damage). The intensity of the heat stimulus was maintained constantly throughout all experiments. Thermal stimuli were delivered four times to each hind paw at 5 to 6 min intervals.

#### Hind paw mechanical withdrawal threshold

The basal pain threshold for mechanical sensitivity was determined by measuring the incidence of foot withdrawal in response to mechanical indentation of the plantar surface of each hind paw. A set of von Frey filaments with a sharp, cylindrical probe and a uniform tip diameter of approximately 0.2 mm (0.04 - 2 g; North Coast medical, Inc.) was used for indetation under a protocol described previously (*62, 63*). In brief, the mouse was placed on a metal mesh floor and covered with a transparent plastic dome (10 × 15 × 15 cm). The animal was allowed to explore and habituate to the chamber for 15 min. Each filament was then applied from underneath the metal mesh floor to the plantar surface of the foot. The duration of each stimulus was 3 s, in the absence of withdrawal, and the interstimulus interval was 10-15 s. The incidence of foot withdrawal was expressed as a percentage of the 10 applications of each stimulus as a function of force. 50% withdraw threshold values were then determined.

#### Rotarod test

The rotarod test was performed using a four-lane Rotarod apparatus (Med Assocaites Inc). The rotarod speed started at 4 Revolutions Per Minute (RPM) and uniformly accelerated to 40 RPM over 5 m. Each mouse was tested 3 times with 5 min intervals.

### Spinal cord slice preparation

Young adult mice of both sexes (7–12 w old) were deeply anesthetized with isoflurane, and lumbosacral laminectomies were performed in an ice-cold chamber. The lumbosacral spinal cord was quickly removed and placed in ice-cold, sucrose-substituted artificial CSF saturated with 95% O_2_ and 5% CO_2_ (sucrose ACSF, in mM: sucrose, 234; KCl, 3.6; CaCl_2_, 2.5; MgCl_2_, 1.2; NaH_2_PO_4_, 1.2; NaHCO_3_, 25; and D-glucose, 12). Transverse lumbar spinal cord slices (300 μm) were prepared in ice-cold sucrose ACSF using a vibrating microtome. The slices were then maintained in a recovery chamber for 30 min or more and studied at room temperature (22–25°C) in regular ACSF equilibrated with 95% O_2_ and 5% CO_2_ (ACSF, in mM: NaCl, 125; KCl, 2.5; CaCl_2_, 2; MgCl_2_, 1, NaH_2_PO_4_, 1.25; NaHCO_3_, 26; and D-glucose, 25, and sucrose added to make 300–320 mOsmol).

### Two-photon imaging

#### Spinal cord in vivo 2-photon imaging

As described previously (*64*), an anesthetic mix of ketamine-xylazine-acepromazine was used to anaesthetize the mouse. Custom-ordered clamps were used to stabilize the spinal column, head, and tail (model STS-A; Narishige), minimizing movement-associated artifacts. After a T12-L1 laminectomy, the lumbar spinal cord was exposed. Under a binocular microscope with 8X to 40X magnification, the dura was cut and removed. A small well of Gelseal (Amersham Biosciences Corp.) was built around the exposed spinal cord to facilitate the maintenance of the tissue in a drop of artificial cerebrospinal fluid (ACSF) and for the immersion of the microscope lens in this solution for *in vivo* imaging. As described above, we used a two-photon microscope (Scientifica Inc., UK) with a Ti:sapphire laser (Mai Tai; Spectra Physics) tuned to 900 nm for *in vivo* imaging. Images of the superficial dorsal horn were obtained by preparing image stacks (167 × 167 μm, 1 μm z steps) collected at a depth of 30-80 μm from the spinal surface before and after formalin injection (or high frequency stimulation). At the end of the experiment, the mice were given an overdose of anesthetic and then killed by cervical dislocation. We used the lowest laser power that would allow for discernment of puncta to avoid phototoxicity. Stacks of images were collapsed to 2D for presentation and quantification using ImageJ software (National Institutes of Health, Bethesda, MD).

#### Spinal cord slice ex vivo 2-photon imaging

Experiments were conducted at room temperature with slices maintained in oxygenated regular ACSF with the same composition as mentioned above in a perfusion chamber at a flow rate of 2 ml/min. GFP-labeled microglia were typically imaged using a two-photon microscope (Scientifica) with a Ti:sapphire laser (Mai Tai; Spectra Physics) tuned to 900 nm with a 40X water-immersion lens (0.8 NA; Olympus). The laser power was maintained at 25mW or below. Microglial were imaged in between 40-100 µm of the spinal cord slice surface. Microglia imaging was limited to 3 hours after slicing to limit microglial activation effects as a confounding variable in data collection.

### Whole-cell patch clamp recordings

For whole-cell, patch clamp recordings, 7–10 week-old adult mice of both sexes were used to perform experiments. Spinal cord slices were prepared 40 min after formalin injection. The whole cell patch-clamp recordings were made from lamina I-III GFP-labeled microglia in voltage-clamp mode. After establishing the whole-cell configuration, microglia were held at −60 mV throughout all experiments. The resistance of a typical patch pipette is 4–6 MΩ. Recording electrodes contained a K^+^-based internal solution composed of (in mM): K-gluconate, 120; NaCl, 5; MgCl_2_, 1; EGTA, 0.5; MgATP, 2; Na_3_GTP, 0.1; HEPES, 10; pH 7.2; and 280 –300 mOsmol. Membrane currents were amplified with an Axopatch 200B amplifier (Multiclamp 700B, Axon Instruments). Signals were filtered at 2 kHz and digitized (DIGIDATA 1440A), stored, and analyzed by pCLAMP (Molecular Devices, Union City, CA).

### Dynamic microglial process motility analysis

Microglial motility analysis was carried out as described previously (*65*). Briefly, 3D image stacks were combined to make 2D projection images for each time point. Next, to account for any x–y tissue drift, 2D projection images were registered using the StackReg plugin. Registered images were smoothened to reduce background noise. To define the cell boundary, an arbitrary threshold was applied uniformly to all images in a given time sequence. To generate difference images, the absolute difference between two sequential thresholded images in a time series was calculated using the ‘‘Difference’’ tool in the ‘‘Image Calculator” of ImageJ. Sequential difference images in a time sequence were used to generate a motility index (MI), which is a percent change in area.

### Process length analysis

A skeleton analysis method was developed to quantify microglia morphology from two-photon images as previously described (*66*) with slight modifications. Two-photon images (20 μm z-stack at 1 μm intervals) were acquired. Two-dimensional (2D) stacked images were made using the ImageJ program. For skeleton analysis, the maximum intensity projection image of the GFP signal was de-speckled to eliminate background noise. The resulting image was converted to a binary image and skeletonized. The Analyze Skeleton plugin (https://imagejdocu.tudor.lu) was then applied to all skeletonized images to collect data on the process length.

### Sholl analysis

For Sholl analysis, Z-stacks were acquired at 1 μm intervals pf from 2-photon live imaging of GFP microglia (20 μm). Consecutive Z-stack images were converted to a maximum intensity projection image using Image J software. Using the Image5D plugin of Image J, Z-stack images were condensed into a maximum intensity projection image over which concentric circles were drawn (concentric circles plugin, Image J), centered on the soma, beginning at 7 μm radii and increasing with every circle. Sholl analysis was manually performed for each cell by counting the number of intersections between microglia branches and each increasing circle to create a Sholl plot.

### Fluorescent immunostaining

Mice were deeply anesthetized with isofluorane (5% in O_2_) and perfused transcardially with 20 ml PBS followed by 30 ml of cold 4% paraformaldehyde (PFA) in PBS containing 1.5% picric acid. The spinal cord was removed and post-fixed with the same 4% PFA solution overnight at 4°C. The samples were then transferred to 15% and then 30% sucrose in PBS overnight. Sample sections (17 µm in thickness) were prepared on gelatin-coated glass slides using a cryostat (Leica). The sections were blocked with 5% goat serum and 0.3% Triton X-100 (Sigma) in TBS buffer for 60 min, and then incubated overnight at 4°C with primary antibody for rabbit anti-Fos polyclonal antiserum (diluted 1:200, Santa Cruz, Cat. #sc-52), rabbit anti-pERK (1:500, Cell Signaling, Cat. #4511), rat anti-mouse F4/80 (1:200, Abd Serotech, Cat. #MCA497), rabbit anti-Iba1 (1:1000, Wako Chemicals, Cat. #019-19741), mouse anti-neuronal nuclei (NeuN) monoclonal antibody (1:200, Chemicon Millipore, Cat. #MAB377), or mouse anti-GFAP (1:500, Cell Signaling, Cat. #12389T). The sections were then incubated for 90 min at RT with secondary antibodies (1:500, Alexa Fluor® 488, 594, Life Technologies). The sections were mounted with Fluoromount-G (SouthernBiotech, Cat. #0100-20) and fluorescent images were obtained. Cell counting and fluorescent signal intensity was quantified using ImageJ software.

### Statistical analysis

All data were expressed as mean ± SEM. Quantification of Iba1 cells, c-fos cells and pERK cells was performed with ImageJ software for statistical analysis using unpaired 2-tailed Student’s t-test. Behavioral data were analyzed using Student’s t-test (2 groups) or Two -way ANOVA followed by post-hoc Bonferroni tests for multiple groups. The criterion for statistical significance was P < 0.05.

## Acknowledgements

This work is supported by National Institute of Health (R01NS088627, R21DE025689, R01NS112144), and National Natural Science Foundation of China (No. 81771183, 81371510). We thank Noriko Goldsmith (Rutgers University) for technical assistance with confocal microscopy. Also, we greatly appreciate all members of Wu lab at Rutgers and Mayo for insightful discussions.

