## Supplemental info and figures for "Spinal Microglia Contribute to Sustained Inflammatory Pain via Amplifying Neuronal Activity"

### SUPPLEMENTARY MATERIALS AND METHODS

### SUPPLEMENTARY FIGURES AND FIGURE LEGENDS

### SUPPLEMENTARY MATERIALS AND METHODS

#### Monocyte flow cytometry

Whole mouse blood was collected and monocytes were separated from erythrocyte and granulocyte on a Ficoll (GE Healthcare) gradient. Separated monocytes were washed with Hank's Balanced Salt Solution (HBSS) and then incubated with 2% goat serum for 10 min, and stained with APC-conjugated CD11b antibody (1:200, Biolegend) for 45 min. Cells were then fixed with 1% PFA for 10 min before flow cytometry. Cells were analyzed on a FACSCalibur cytometer (Becton Dickinson) using the CellQuest software (Becton Dickinson).

#### Microglia cell area and process index analyses

Process index analysis was done by two independent observers who manually counted the number of primary processes emanating from microglial somata. Observers were blinded to control or formalin-injected conditions. Data from the two observers were averaged. Microglial morphology analysis was performed using ImageJ. An automated threshold was set on each image and a particle analysis was performed on thresholded images. Microglia cell area was averaged and normalized to 100% for control microglia and compared with microglia after formalin-induced acute pain.

#### High-frequency stimulation

The high-frequency stimulation operation was performed under anesthesia using an anesthetic mix of ketamine-xylazine-acepromazine. The left sciatic nerve was carefully exposed at mid-thigh level under aseptic conditions. A pair of silver electrode hooks was inserted under the nerve. When stimulating the nerve, the hindleg and pelvis were fixed tightly to avoid contact with muscle caused by muscle contraction. Briefly, 4 trains of 100Hz stimulation (0.5 ms rectangular pulses at 20V) were applied for a 2-sec duration, with a 10-sec interval between trains. After stimulation, the muscle and skin were sutured in layers. The sciatic nerve of the sham-operated mice was identically exposed and manipulated but was not stimulated.

SUPPLEMENTARY FIGURES AND FIGURE LEGENDS

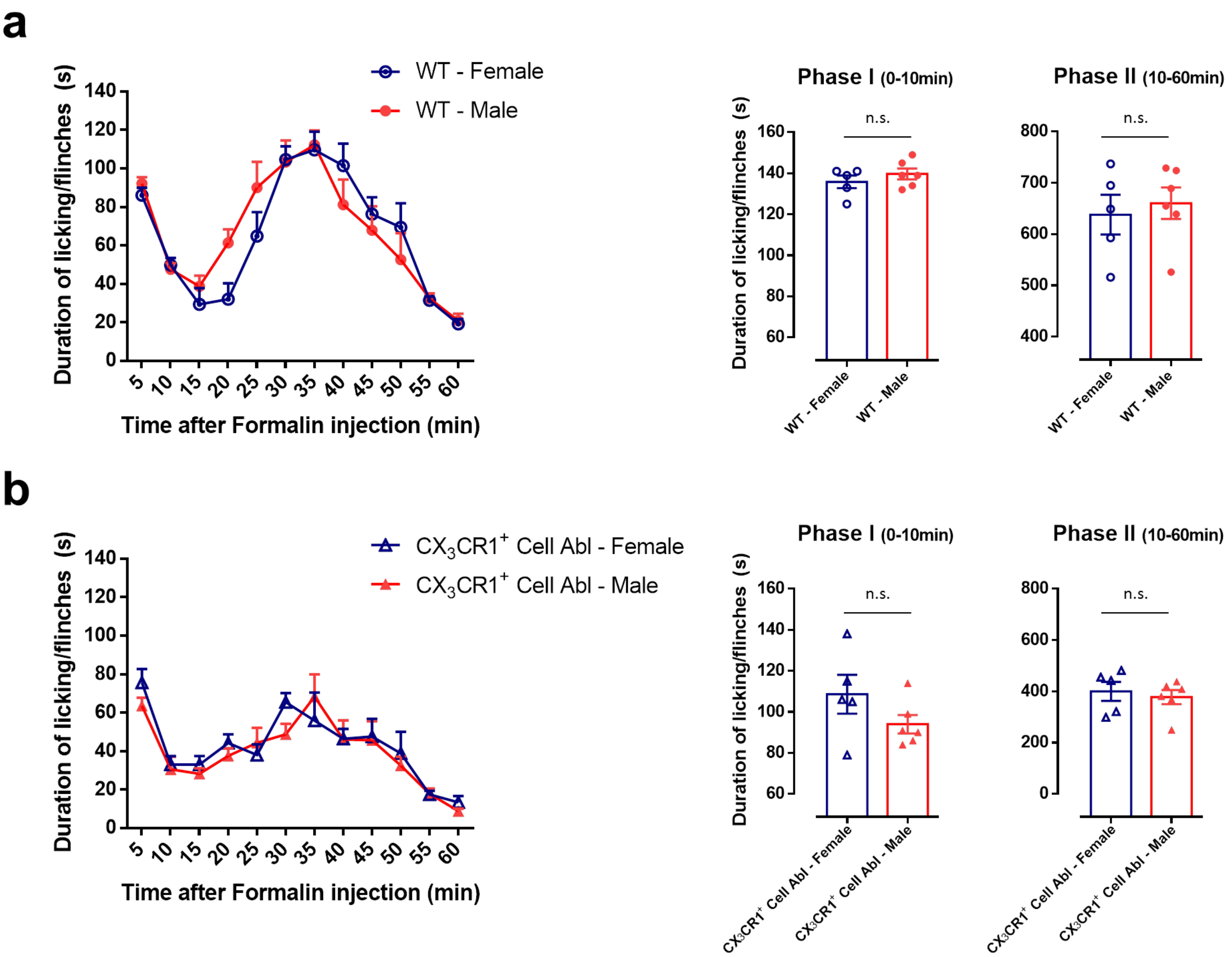


**Figure S1. There is no sex differences in phase I or phase II inflammatory pain responses**

**(a)** Time course (0–60 min) of formalin-induced spontaneous pain behavior (licking/flinching) in wild type male and female mice, as measured every 5 min. Histogram representing formalin-induced Phase-I (1–10 min) and Phase-II inflammatory pain responses (10–60 min). No sex differences were found in both phase I and phase II responses in wild type males and females. Data are presented as mean ± SEM; n.s., no significance, compared to wild type female mice, unpaired 2-tailed Student's t test, n = 5–6 mice/group.

**(b)** Time course (0–60 min) of formalin-induced spontaneous pain behavior (licking/flinching) in $CX_3CR1^+$ cell ablation male and female mice, as measured in every 5 min. Histogram representing formalin-induced Phase-I (1–10 min) and Phase-II inflammatory pain responses (10–60 min) in $CX_3CR1^+$ cell ablation male and female mice. No sex differences were found in both phase I and phase II responses in $CX_3CR1^+$ cell ablation males and females. Data are presented as mean ± SEM; n.s., no significance, compared to $CX_3CR1^+$ cell ablation female mice, unpaired 2-tailed Student's t test, n = 5–6 mice/group.

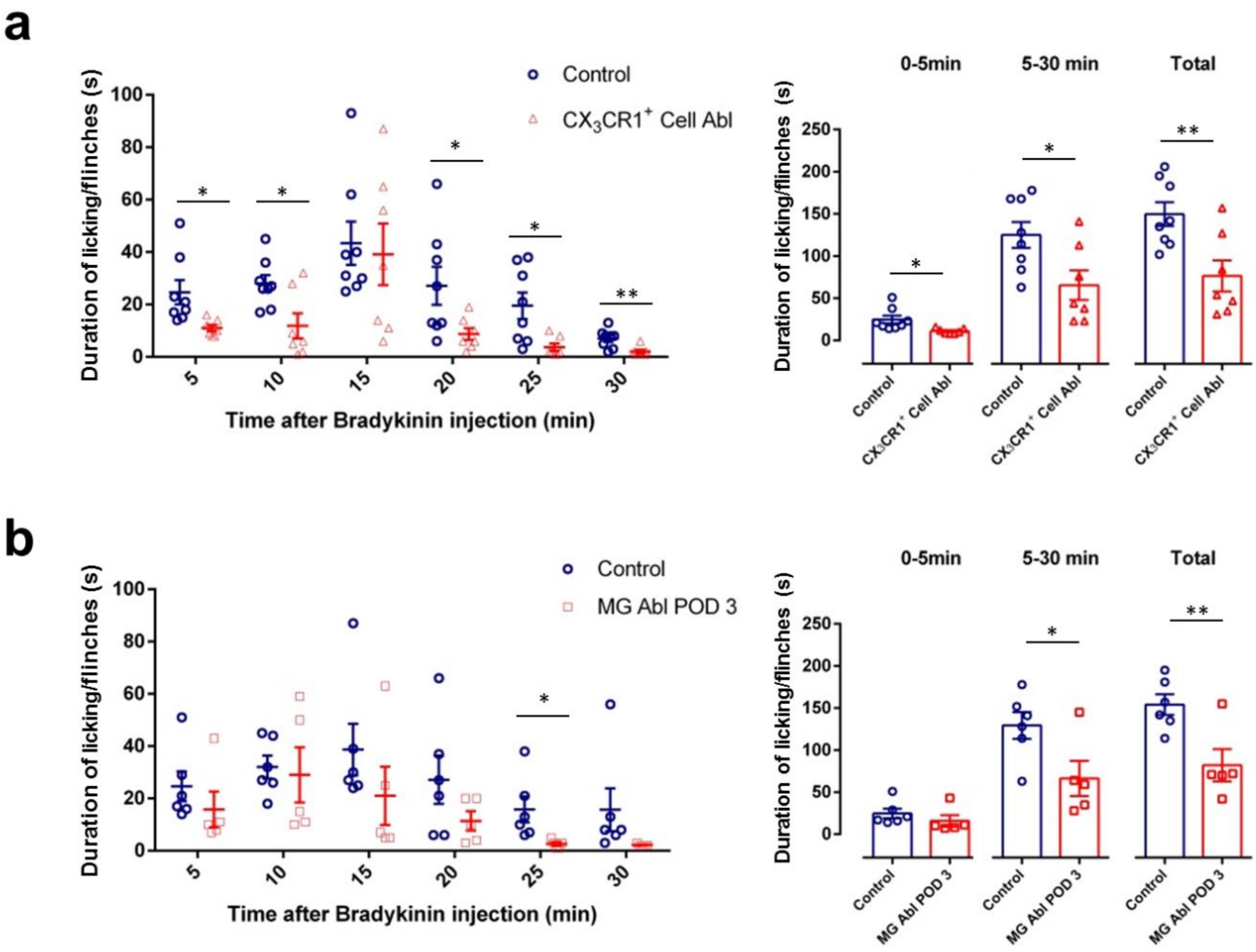


**Figure S2. Bradykinin-induced inflammatory pain is reduced in $CX_3CR1$+ cell ablation mice and microglia ablation mice.**

**(a)** Time course (0–30 min) of bradykinin-induced spontaneous pain behavior (licking/flinching) in control and $CX_3CR1^+$ cell ablation mice, as measured every 5 min. Histogram representing bradykinin-induced inflammatory pain responses within 0–5 min, 5–30 min and 0-30 min in control and $CX_3CR1^+$ cell ablation mice. Bradykinin-induced inflammatory pain is reduced across both observation periods in $CX_3CR1^+$ cell ablation mice. Data are presented as mean ± SEM; *$P < 0.05$, **$P < 0.01$, compared to control mice, unpaired 2-tailed Student's t test, n = 7–8 mice/group.

**(b)** Time course (0–30 min) of bradykinin-induced spontaneous pain behavior (licking/flinching) in control and microglia ablation POD3 mice, as measured every 5 min. Histogram representing bradykinin-induced inflammatory pain responses within 0–5 min, 5–30 min and 0-30 min in control and microglia ablation POD3 mice. Bradykinin-induced inflammatory pain is reduced in 5–30 min and 0-30 min in microglia ablation POD3 mice. Data are presented as mean ± SEM; *P < 0.05, **P < 0.01, compared to control mice, unpaired 2-tailed Student's t test, n = 5–6 mice/group.

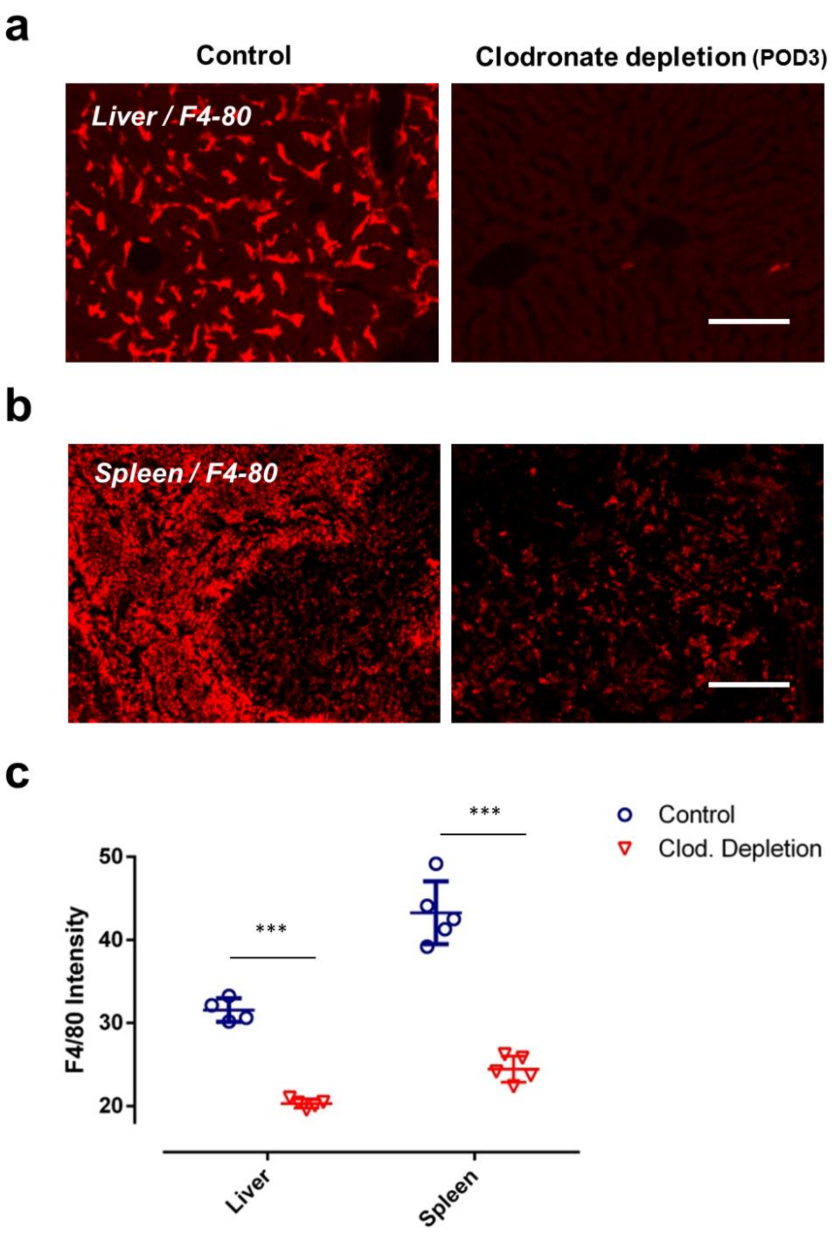


**Figure S3. Macrophages in liver and spleen are largely depleted after clodronate treatment.**

**(a-b)** Representative images of F4/80-positive macrophages in liver **(a)** and spleen **(b)** from control and clodronate depletion mice. Scale bar = 400 μm. **(c)** Quantitative data showing the average density of F4/80- positive cells in liver and spleen from control and clodronate depletion mice. Data are presented as mean ± SEM. ***p < 0.001, compared to control mice, unpaired 2-tailed Student's t test, n = 4–5 mice/group.

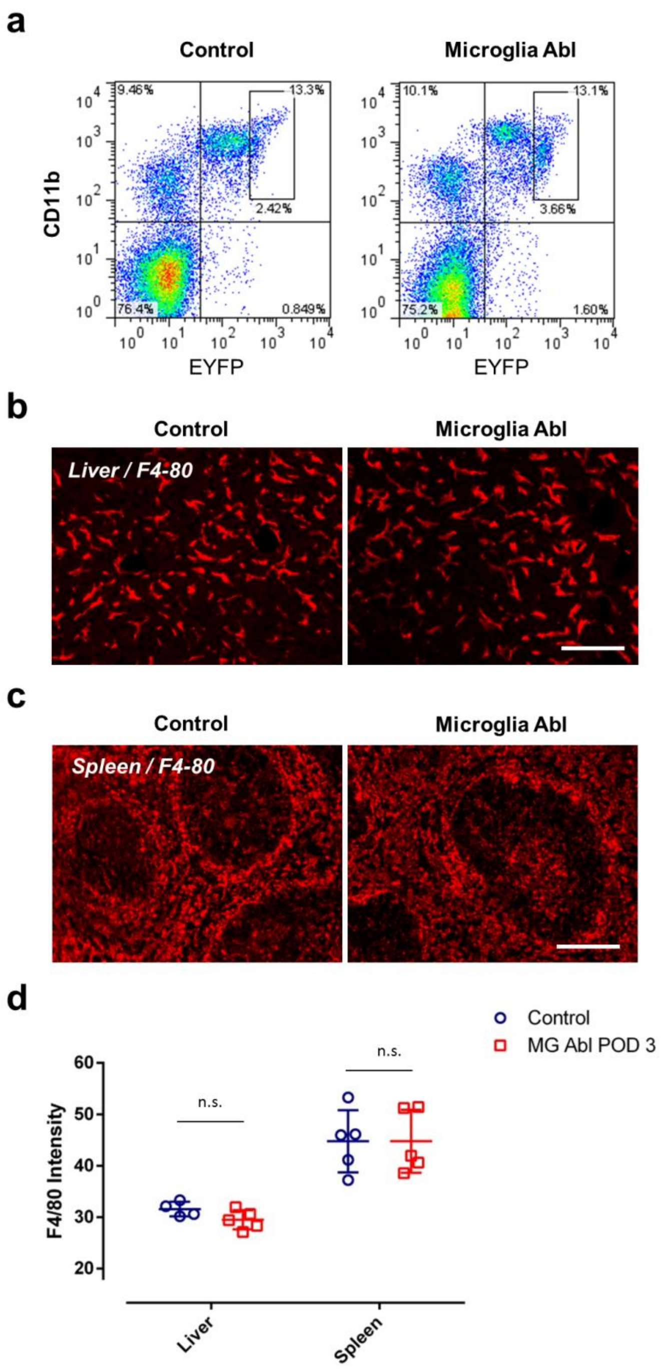


**Figure S4. Macrophages in the blood, liver and spleen in microglia ablation mice.** **(a)** Blood monocyte cytometry showing a preserved CD11b+$CX_3CR1$+ cell population in microglia ablation mice compared to control mice. $CX_3CR1$ expression was detected through EYFP fluorescence ($CX_3CR1$-$^{CreER\text{-}EYFP}$). **(b-d)** F4/80-positive macrophages in liver **(b)** and spleen **(c)** remain unaffected in microglia ablation mice. Scale bar = 400 μm. Quantitative data **(d)** showing the average density of F4/80- positive cells in control and microglia ablation mice. Data are presented as mean ± SEM. n.s., no significance, compared to control mice, unpaired 2-tailed Student's t test, n = 4–5 mice/group.

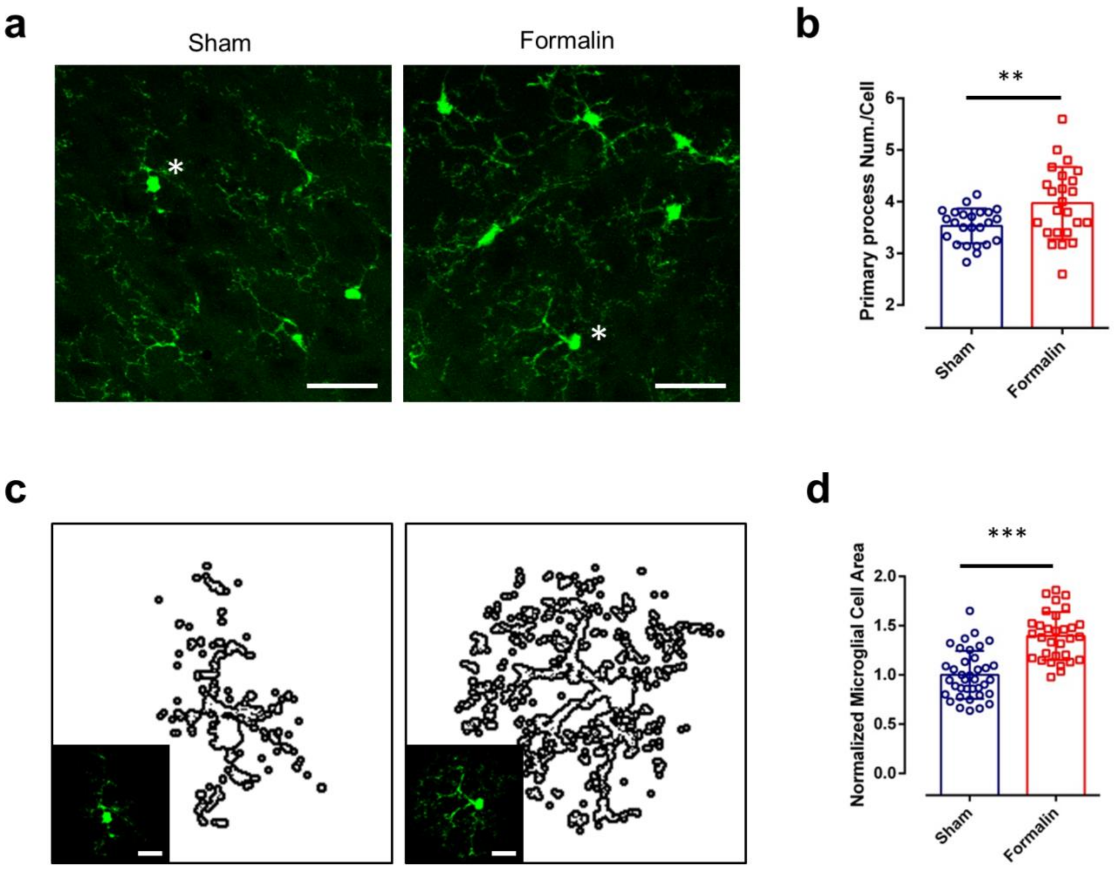


**Figure S5. Intraplantar formalin injection alters microglial morphology in fixed sections.**

**(a)** Representative images of microglia in spinal cord dorsal horn Lamina I-III from sham and formalin-treated mice 1 hr post injection. Scale bar = 30 μm.

**(b)** Quantitative data showing intraplantar formalin injection increases primary microglial branch number. Data are presented as mean ± SEM. **p < 0.01, compared to sham mice, unpaired 2-tailed Student's t test, n = 24 microglia from 4 mice per group.

**(c)** Representative images of outlined microglia in representative slices (Asterisks in Figure S5a) from sham and formalin-treated mice used to determine microglial area. Scale bar = 15 μm.

**(d)** Quantitative normalized area showing intraplantar formalin injection increases microglial cell area. Data are presented as mean ± SEM. ***p < 0.001, compared to sham, unpaired 2-tailed Student's t test, n = 31–37 microglia from 4 mice per group.

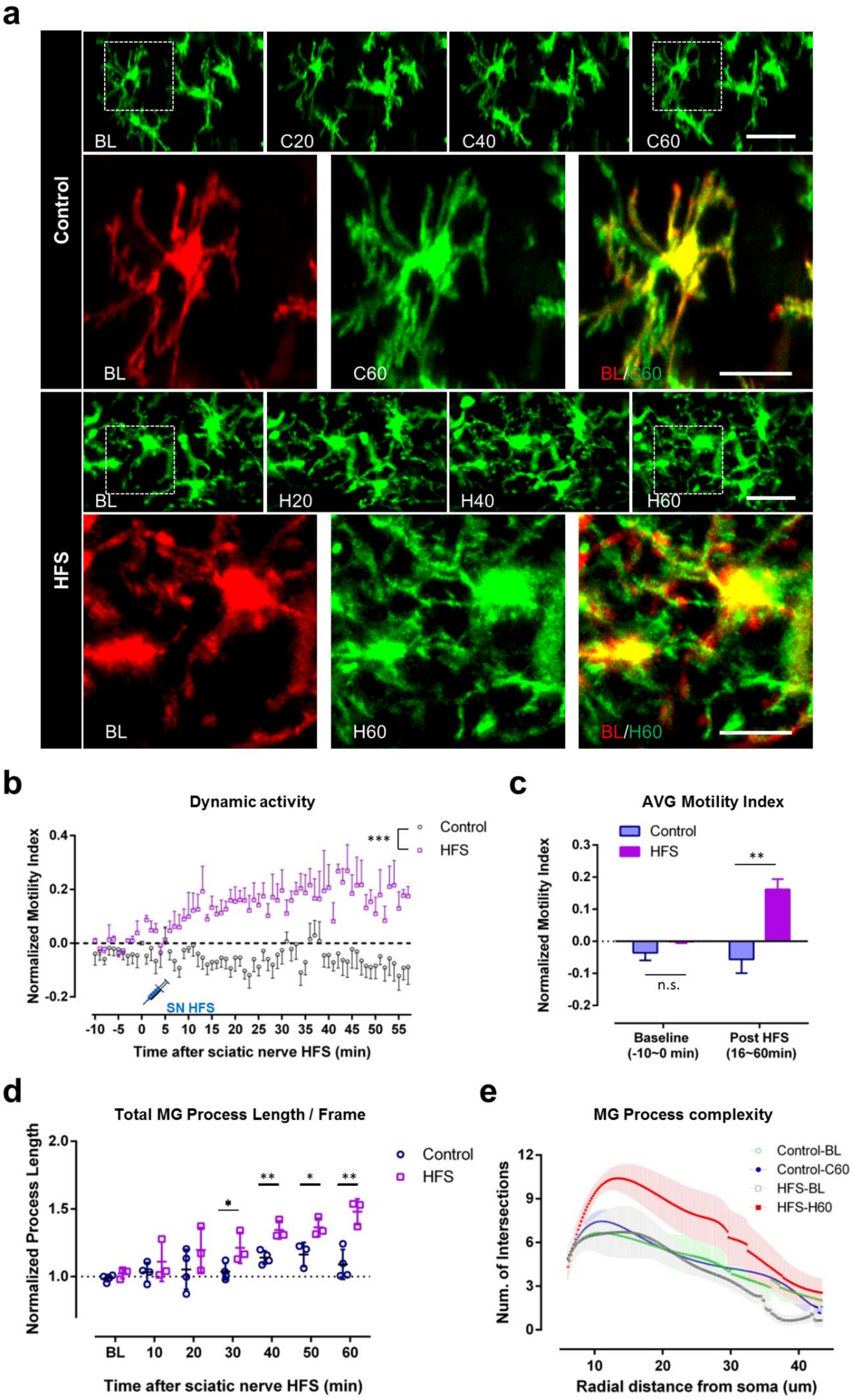
a
Control
BL
C20
C40
C60
BL
C60
BL/C60
HFS
BL
H20
H40
H60
BL
H60
BL/H60
b
Dynamic activity
Normalized Motility Index
***
Control
HFS
SN HFS
Time after sciatic nerve HFS (min)
c
AVG Motility Index
Control
HFS
Normalized Motility Index
**
n.s.
Baseline
(-10~0 min)
Post HFS
(16~60min)
d
Total MG Process Length / Frame
Normalized Process Length
Control
HFS
BL
Time after sciatic nerve HFS (min)
e
MG Process complexity
Num. of Intersections
Control-BL
Control-C60
HFS-BL
HFS-H60
Radial distance from soma (um)

**Figure S6. Increased complexity and dynamics of spinal microglial processes after sciatic nerve stimulation.**

**(a)** Representative *in vivo* z-stack images of L4-5 ipsilateral spinal microglia taken before (–10 min, BL) and after either high frequency stimulation (HFS) or sham stimulation of the sciatic nerve. Microglial dynamics were studied 20, 40, and 60 minutes after each condition (Sham: C20, C40, C60; HFS: H20, H40, H60). Higher magnification images of a microglial cell before (–10 min; red) and 60 min after HFS or control conditions (green) are shown individually and in a merged image to display changes in process dynamics. Scale bars represent 40 μm and 20 μm**.**

**(b)** Quantitative data showing dynamics of spinal microglial processes, represented as the normalized motility index, in control and HFS stimulation mice. Data are shown as mean ± SEM. ***P < 0.001, compared to control. Two-way ANOVA. n = 4 for control and n = 3 for HFS stimulation mice.

**(c)** Quantitative data showing average motility index of spinal microglial processes in control and sciatic nerve HFS stimulation groups at different time periods (-10-0 min before and 16-60 min after sciatic nerve HFS stimulation. Data are shown as mean ± SEM. **P < 0.01, compared to Control. Unpaired 2-tailed Student's t test. n = 4 for control and n = 3 for sciatic nerve HFS stimulation.

**(d)** Quantitative data showing spinal microglial processes length, represented as normalized process length, in control and s HFS stimulation at different time points. Process lengths were analyzed every 10 min. Data are shown as mean ± SEM. *P < 0.05, **P < 0.01, compared to control. Unpaired 2-tailed Student's t test. n = 3-4/group.

**(e)** Sholl analysis plot indicating microglial branch complexity in control and HFS stimulation groups. n = 6-7/group. Data are shown as mean ± SEM.
